# Scaled and Efficient Derivation of Loss of Function Alleles in Risk Genes for Neurodevelopmental and Psychiatric Disorders in Human iPSC

**DOI:** 10.1101/2024.03.18.585542

**Authors:** Hanwen Zhang, Lilia Peyton, Ada McCarroll, Sol Díaz de León Guerrerro, Siwei Zhang, Prarthana Gowda, David Sirkin, Mahmoud El Achwah, Alexandra Duhe, Whitney G. Wood, Brandon Jamison, Gregory Tracy, Rebecca Pollak, Ronald P. Hart, Carlos N. Pato, Jennifer G. Mulle, Alan R. Sanders, Zhiping P. Pang, Jubao Duan

## Abstract

Translating genetic findings for neurodevelopmental and psychiatric disorders (NPD) into actionable disease biology would benefit from large-scale and unbiased functional studies of NPD genes. Leveraging the cytosine base editing (CBE) system, here we developed a pipeline for clonal loss-of-function (LoF) allele mutagenesis in human induced pluripotent stem cells (hiPSCs) by introducing premature stop-codons (iSTOP) that lead to mRNA nonsense-mediated-decay (NMD) or protein truncation. We tested the pipeline for 23 NPD genes on 3 hiPSC lines and achieved highly reproducible, efficient iSTOP editing in 22 NPD genes. Using RNAseq, we confirmed their pluripotency, absence of chromosomal abnormalities, and NMD. Interestingly, for three schizophrenia risk genes (*SETD1A, TRIO*, *CUL1*), despite the high efficiency of base editing, we only obtained heterozygous LoF alleles, suggesting their essential roles for cell growth. We replicated the reported neural phenotypes of *SHANK3*-haploinsufficiency and found *CUL1*-LoF reduced neurite branches and synaptic puncta density. This iSTOP pipeline enables a scaled and efficient LoF mutagenesis of NPD genes, yielding an invaluable shareable resource.

## Introduction

In the past decade, genome-wide association studies (GWAS) (Consortium., 2014; Grove et al., 2019; Howard et al., 2019; Meng et al., 2024; Mullins et al., 2021; Purcell et al., 2009; Ripke et al., 2013; Ripke et al., 2011; Shi et al., 2009; Stahl et al., 2019; Stefansson et al., 2009; Trubetskoy et al., 2022; Wray et al., 2018) and whole-exome sequencing (WES) studies (Satterstrom et al., 2020; Singh et al., 2022b) on neurodevelopmental and psychiatric disorders (NPD) such as schizophrenia (SZ), autism spectrum disorder (ASD), bipolar disorder, and major depression, have identified a growing number of risk genes. However, translating these exciting genetic discoveries into translational actionable biology has been impeded by our limited knowledge of gene function and related disease mechanisms. A bottleneck in the field is that genes are often studied individually, slowing the progress and posing potential bias in functional interpretation. To overcome such limitations, the NIMH (National Institute of Mental Health)-initiated SSPsyGene (Scalable and Systematic Neurobiology of Psychiatric and Neurodevelopmental Disorder Risk Genes) Consortium (sspsygene.ucsc.edu) aims to functionally characterize the contribution of 150-250 NPD genes. The selected NPD genes mostly have disease-associated rare protein-truncating variants (PTVs) that likely cause gene loss-of-function (LoF) (Palmer et al., 2022; Satterstrom *et al*., 2020; Singh *et al*., 2022b) and have strong effect sizes (sspsygene.ucsc.edu/resources), which will help interpret their individual biological relevance and determine any convergent or divergent biology across disorders. Large-scale, unbiased, and parallel study of these NPD genes in disease-relevant model systems will substantially deepen our understanding of the pathophysiology of NPD.

Human induced pluripotent stem cells (hiPSC) and their derived neural cells empowered by CRISPR-mediated gene editing provide promising cellular models for studying NPD genes (De Los Angeles et al., 2021; Duan, 2023; Michael Deans and Brennand, 2021; Muhtaseb and Duan, 2022; Wang et al., 2020; Wen et al., 2016) and for scaling up the assay. A “cell village” approach (Wells et al., 2023) enables the co-culture of tens to hundreds of hiPSC lines in a dish and to differentiate them into neurons together, followed by assaying a specific cellular phenotype and genetically inferring individual cell identity. With a similar “cell village” approach, pooled screening of isogenic CRISPR/Cas9-edited hiPSC lines carrying 30 different ASD mutations in 2D neural culture (Cederquist et al., 2020) or combining parallel genetic perturbations of 36 high-risk ASD genes with single-cell transcriptomic readout in mosaic cerebral organoids (Li et al., 2023), has been used to identify LoF phenotypes at both cellular and molecular levels. Such pooled screening of LoF allelic effects may be further scaled up by increasing the number of targeted genes to hundreds or thousands using CRISPRi (Holtzman and Gersbach, 2018) or CRISPRoff (Nunez et al., 2021). While an invaluable approach, the pooled CRISPR screening in hiPSC-derived neural models is limited by cell line-specific or LoF allele-specific unequal cellular growth, possible non-autonomous effects, and the restrictive phenotypes amenable for screening.

A scaled and efficient approach to generate clonal individual hiPSC lines carrying LoF alleles for a large number of NPD genes would complement the pooled CRISPR screening, providing a sharable resource that would enable cellular phenotyping at an unmatched scale. CRISPR/Cas9 editing can be used to systematically create small DNA insertions or deletions (indels) or exon deletions in protein-coding regions through non-homologous end joining (NHEJ) repair of double-strand breaks (DSBs) (Ran et al., 2013), resulting in protein-truncating mutations. Alternatively, LoF mutation can be generated by using CRISPR-based cytosine base editors (CBE) to introduce premature protein stop-codons (i.e., nonsense mutations; an iSTOP approach) that lead to mRNA nonsense-mediated-decay (NMD) and/or protein truncates (Billon et al., 2017; Cuella-Martin et al., 2021; Hanna et al., 2021; Xu et al., 2021). Compared to the traditional CRISPR/Cas9 gene editing system, the CBE editor makes “C” to “T” changes in DNAs without creating cell-toxic DSBs as the Cas9 nuclease does (Ran *et al*., 2013) and with minimized potential off-target DNA editing (Billon *et al*., 2017; Cuella-Martin *et al*., 2021; Hanna *et al*., 2021; Xu *et al*., 2021). Furthermore, compared to the CRISPR/Cas9 editing-induced small indels that may or may not disrupt a protein sequence reading frame, a CBE base editor can precisely introduce a premature stop codon, which makes the clonal LoF allelic confirmation more straightforward and cost-effective in a scaled LoF mutagenesis workflow. Finally, the CBE-engineered premature stop-codon mutations are more reminiscent of the rare patient-specific PTVs or LoF mutations associated with NPD (Satterstrom *et al*., 2020; Singh et al., 2022a). Despite some of its advantages over the CRISPR/Cas9 editing system, the CBE editor often has much lower editing efficiency in hiPSC. Although the DNA base editing reporter gene system has been developed to enrich the edited cells, thereby increasing the base editing efficiency of a target gene (Standage-Beier et al., 2019), the use of a CBE editor in editing hiPSC lines has been scarce, and its usefulness in developing a scaled and efficient clonal LoF mutagenesis in hiPSC has not been tested.

As part of the SSPsyGene Consortium, our Assay and Data Generation Center (ADGC) for the Model of iPSC-derived Neurons for NPD (MiNND) aims to employ the CBE editor-based iSTOP approach to generate isogenic hiPSC lines carrying LoF alleles for about 150-200 NPD genes on multiple donor genetic backgrounds. Here, leveraging an improved reporter gene editing enrichment system that can substantially increase the CBE iSTOP editing efficiency in hiPSC, we established a semi-automated pipeline for parallel and efficient clonal LoF mutagenesis of a large number of genes. We tested the workflow on 23 NPD genes with 3 donor hiPSC lines (KOLF2.2J, CW20107, MGS_CD14). We obtained high and reproducible iSTOP editing efficiency across all three hiPSC lines. We systematically characterized the engineered isogenic iSTOP hiPSC lines for pluripotency, karyotyping, neuron differentiation capacity, and the expected NMD and LoF.

## Results

### The CBEmax DNA base-editing enriching system substantially increases “C” to “T” editing in hiPSC

A key for generating LoF alleles by using a CBE editor to introduce premature stop codons (C to T changes; i.e., iSTOP approach) (Billon *et al*., 2017; Popp and Maquat, 2016) on a large scale is to have sufficiently high gene editing efficiency. Although DNA base editors have high SNP editing efficiency (>50%) in some commonly used cell lines such as HEK293 (Rees and Liu, 2018), hiPSCs are less tested. We opted to employ a base editing reporter gene system to enrich the gene-edited cells (Standage-Beier *et al*., 2019), thereby increasing iSTOP editing efficiency of a target gene in selected cells. In this CBE editing enriching system (CBEmax_Enrich), a blue fluorescent protein (BFP) reporter on the reporter plasmid pEF-BFP will turn into a functional EGFP reporter when it is edited from CAC (H66) to TAC (Y66) in cells co-transfected with pEF-AncBE4max and sgRNAs (Figure 1A). We first individually tested the two iSTOP sgRNAs (Table S1) that target the *apolipoprotein E* (*APOE*) gene in HEK293 cells. For each sgRNA, we found a substantial increase in the target gene editing efficiency (C to T) in fluorescence-activated cell sorting (FACS)-sorted GFP+ cells compared to the transfected BFP+ cells (from 42% to 86% and from 29% to 81%, respectively) (Figure 1B). Because a regular CBE editor such as AncBE4max has a preference for the protospacer adjacent motif (PAM), which would limit the target regions for designing sgRNAs, we also constructed a “PAM-less” CBE editor that contains a SpRY nuclease (Walton et al., 2020) (plenti-CBE3.9max-SpRY; Figure S1A) that would offer more flexibility for designing sgRNAs in our scaled iSTOP editing (although the PAM-less system was not used in the currently reported batch of LoF mutagenesis). We then tested its gene editing performance in the reporter gene editing enrichment system. Despite a smaller fold-increase of editing efficiency than AncBE4max (from 7% to 33%), our “PAM-less” CBE editor did show a robust increase of C to T editing efficiency (from 7% in the transfected cells to 18% in the enriched GFP+ cells) (Figure S1B).

**Figure 1.**
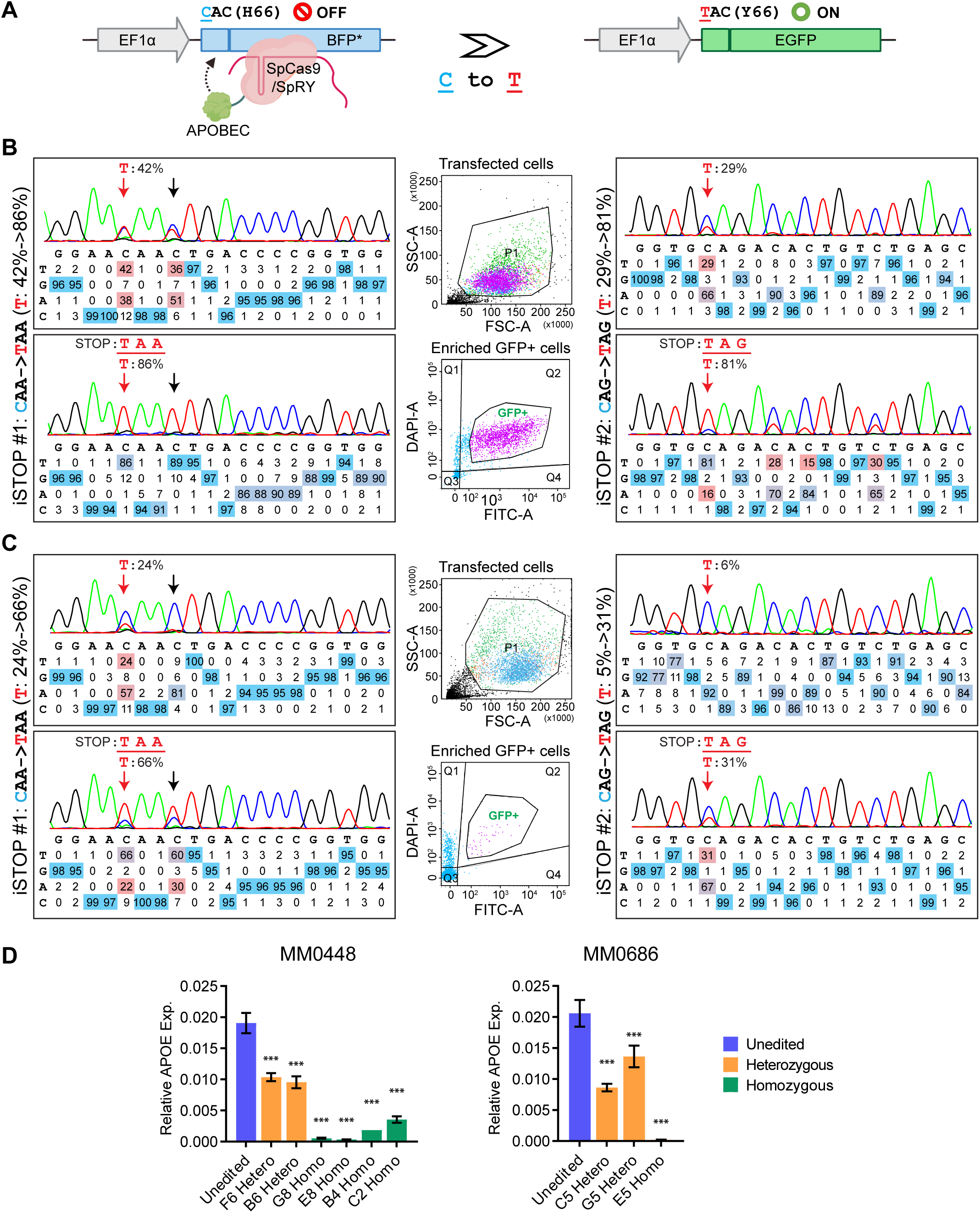
Improved iSTOP base editing efficiency by enriching cells with the reporter gene edited. (A) BFP cassette of the CBEmax_enrich reporter vector. C to T change turns BFP to EGFP in cells undergoing base editing. (B) High C to T editing efficiency of two iSTOP sgRNAs (iSTOP#1 on the left and iSTOP#2 on the right) in HEK293 cells upon enrichment. Middle panels show the cell gating patterns of the dissociated single cells (transfected; upper) and the editing-enriched GFP+ cells (lower panel). (C) Improved C to T editing efficiency in hiPSC for the same iSTOP sgRNAs as in (B). (D) Target gene (APOE) expression knockdown in hiPSC lines homozygous or heterozygous for the T allele after editing using the iSTOP1 and iSTOP2 sgRNAs. Two donor lines are shown and the mRNA expression in different hiPSC clones were quantified by qPCR. The relative expression value was normalized to GAPDH expression. Two-tailed Student’s *t*-test with unequal variance was used for comparison between unedited and edited clones, n = 3 biological replicates. * *P* <0.05, ** *P* < 0.01, *** *P* < 0.001.

We next tested for the iSTOP editing efficiency in two hiPSC cell lines and whether the introduced iSTOP codons led to the expected NMD (i.e., LoF). For both iSTOP sgRNAs, we observed a robust increase, although to a less extent than in HEK293, of the target gene editing efficiency in FACS-sorted GFP+ cells compared to the transfected BFP+ cells (from 24% to 66% and from 5% to 31%, respectively) (Figure 1C). More importantly, as expected from the iSTOP-mediated NMD of mRNAs, we found 86% and 98% of APOE expression reduction in hiPSC clones homozygous for iSTOP1 and iSTOP2, respectively, and ∼50% expression reduction in hiPSC clones heterozygous for iSTOP mutations (Figure 1D).

Taken together, these results show the CBEmax_Enrich system can significantly increase the iSTOP editing efficiency, which enables us to generate LoF alleles on a large scale by introducing premature stop codons.

### A scalable workflow for efficiently deriving clonal LoF alleles in hiPSC using CBEmax_enrich

Our goal is to develop an efficient pipeline that involves single hiPSC cell sorting for deriving clonal LoF alleles of hundreds of NPD genes in multiple hiPSC lines. To achieve this goal, besides the increase of iSTOP editing efficiency by the CBEmax_Enrich system, another key factor is to obtain a relatively high single hiPSC clonal survival rate after FACS of the enriched GFP+ cells (Figures 1B, 1C). It has been recently shown that the CEPT small molecular cocktail can increase single hiPSC cloning efficiency after cell sorting when compared to Rock inhibitor (Y-27632; ROCK-I) (Tristan et al., 2023). We thus tested the performance of CEPT by treating the hiPSC with CEPT both during CBEmax_Enrich transfection (for iSTOP sgRNAs of 4 genes) and the FACS-sorting of single cells into 96-well plates 48hr post transfection. However, we observed a very low single hiPSC clonal survivability (∼5%) despite a high editing efficiency (∼70%) (Figures S2A, S2B). Interestingly, we found that combining our routine ROCK-I treatment of hiPSC at transfection with CEPT treatment during 48hr post-transfection cell sorting gave us a much higher single hiPSC clonal survivability (∼27%), and even higher survivability (∼35%) when we sorted cells 72 hr post-transfection, while maintaining the high gene editing efficiency (Figures S2A, S2B).

After these optimizations to achieve high gene editing efficiency and single hiPSC clonal survivability, we designed a semi-automated pipeline for deriving clonal LoF alleles in hiPSC for 23 NPD genes for each batch (Figure 2A). Briefly, the CBEmax_enrich vector, together with reporter BFP plasmid and gRNA vector carrying the reporter sgRNA and a targeting sgRNA, were transiently transfected into hiPSC in a 24-well plate, each well with one of the 23 targeted LoF mutations or a non-transfection (gRNA)-control (NTC) for 1 donor hiPSC line. We then sorted out single cells that are GFP+ (thus enriched for base editing) and distributed 96 single cells per gene/LoF in a 96-well plate. A handful of single hiPSC colonies from each 96-well plate were further subject to Sanger sequencing to verify the C to T changes (LoF allele) introduced into each NPD gene, and 2-3 hiPSC clones, preferably homozygous for a LoF allele, were banked. The selected hiPSC clones were also subject to RNA-seq to confirm the absence of chromosomal abnormality by eSNP-Karyotyping and pluripotency test (see below). With this pipeline, we have generated LoF alleles, mostly homozygous, for 22 of the 23 selected SSPsyGene Consortium-prioritized NPD genes (no editing found for *HERC1, Table S3*), including 9 (*ARID1B, CACNA1G, CHD8, DLL1, GABRA1, KMT2C, SCN2A, SHANK3, SMARCC2*) out of 10 “capstone genes” for which we could design sgRNAs, on 3 donor lines of European ancestry (KOLF2.2J, CW20107, MGS_CD14) (Figure 2B; Table S2). Our MiNND project within the SSPsyGene consortium is to produce LoF alleles for about 150-200 NPD genes on 6 different hiPSC lines, including two of African Ancestry (Table S2). The derivation of a large number of iSTOP LoF alleles enables us to systematically evaluate the performance of iSTOP base editing on hiPSC and its efficiency in leading to LoF (see below).

**Figure 2.**
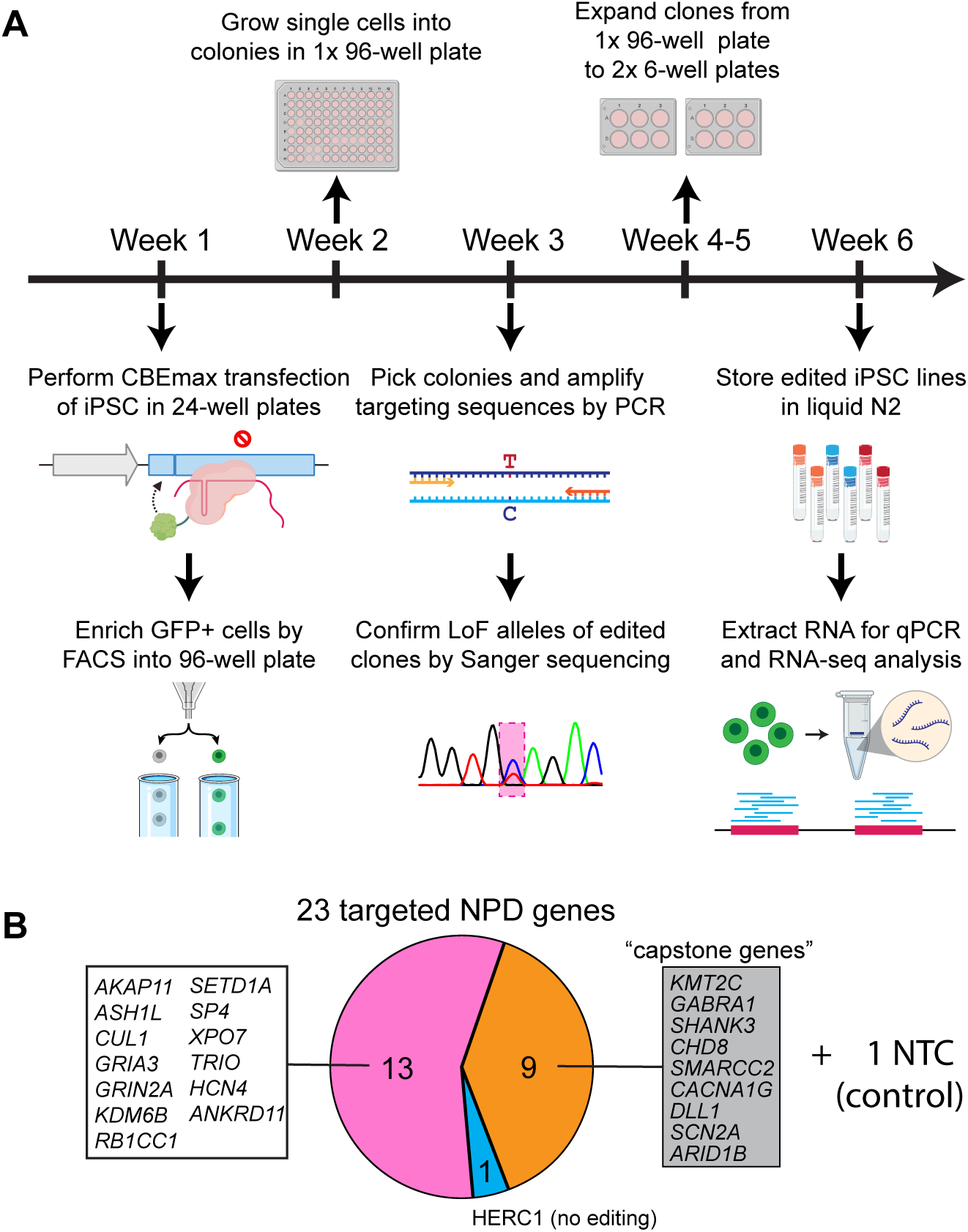
The efficient iSTOP base editing pipeline to introduce LoF alleles in large scale. (A) The workflow that enables the iSTOP editing in batches of 24 (23 target genes + 1 control). The hiPSC transfection was performed on 24-well plate, followed by single hiPSC sorting, single clone expansion on 96-well plate, clonal sequencing confirmation and hiPSC banking. (B) Genes and hiPSC lines used in the current study. Nine prioritized genes (capstone genes) by SSPsyGene consortium are listed in the grey box. Other genes are those NPD risk genes selected by the SSPsyGene consortium to have strongest disease associations and highest priorities for creating LoF alleles.

### iSTOP CBE base editing in hiPSC is efficient and reproducible in different hiPSC lines

The performance of the CBE editor in hiPSC, especially in the context of the iSTOP design and with reporter gene editing enrichment, has not been systematically evaluated previously. With data from the iSTOP base editing of 23 genes across 3 donor hiPSC lines (Figure 2), we found on average the post-editing single-cell clonal viability to be 35-47% (Figure 3A) and the reporter gene editing efficiency to be 31-50% (Figure S2C). After reporter gene editing enrichment, the average target gene editing efficiency was ∼60%, with a strong correlation among different cell lines (Pearson R=0.91-0.95) (Figures 3B, S3A, S3B), demonstrating the highly efficient and reproducible iSTOP CBE editing across all three hiPSC lines. About half of the genes showed editing efficiency higher than 90% and only 5 genes with editing efficiency less than 10% (including the one without editing) (Figure 3B). Despite the robust increase of target gene editing efficiency after reporter gene editing enrichment (Figure 1), there was a weak correlation between reporter gene editing efficiency and target gene editing efficiency (Figure S3C), suggesting target gene editing efficiency is predominately determined by gene-specific sgRNA performance.

**Figure 3.**
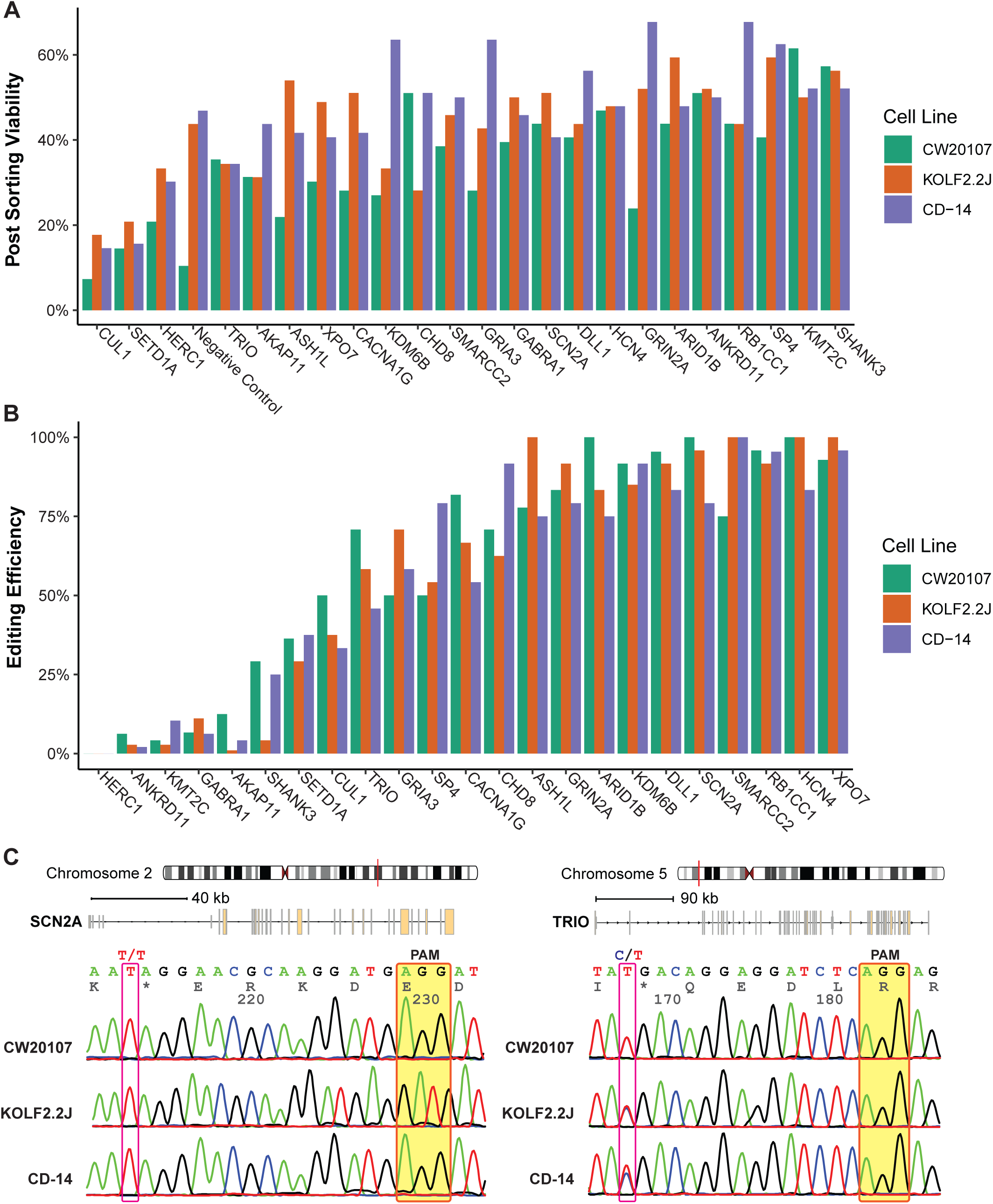
High and reproducible C to T base editing efficiencies across genes and hiPSC lines. (A) High rate of single hiPSC clonal survivability after post-transfection cell sorting. (B) High iSTOP LoF allele editing efficiency and reproducibility. The genotypes were confirmed by Sanger sequencing for the selected individual hiPSC clones from each gene editing. (C) Examples of Sanger sequencing traces to confirm LoF alleles of two genes, *SCN2A* (left) and *TRIO* (right), in all three hiPSC lines. The shown sequencing traces are near the iSTOP-sgRNA region, with PAM sequence highlighted in transparent yellow boxes and the genotype of the LoF mutation site marked in red line box.

Overall, we obtained clonal hiPSC lines carrying putative LoF alleles for 22 targeted genes (no editing found for *HERC1*), of which 15 are homozygous (Figure 3B). We found that although the genes with heterozygous LoF alleles tended to have low editing efficiency (<10%) (*ANKRD11*, *KMT2C*, *GABRA1*, *AKAP11*), some had high editing efficiency (*SETD1A* with 29-38%, *TRIO* with 46-71%, *CUL1* with 33-50%) (Figure 3B), suggesting that for some NPD genes homozygous LoF alleles may have deleterious effects on hiPSC survival or growth. It is noteworthy that all three genes (*SETD1A*, *TRIO*, *CUL1*) with only heterozygous LoF clones, despite their relatively high editing efficiency, are top-ranking SZ risk genes found by the SZ Exome Sequencing Meta-Analysis (SCHEMA) consortium to have rare and highly penetrant SZ-associated PTVs (Singh *et al*., 2022b). Of these genes, *TRIO* was found to initially have homozygous hiPSC clones grown in the post sorting 96-well plate, however, only heterozygous clones (Figure 3C) were found to show sustained normal hiPSC growth, which is consistent with the known necessary role of *TRIO* for cell migration and growth (Deinhardt et al., 2011; Seipel et al., 1999).

### The CBE-edited iSTOP hiPSC clones are pluripotent and have minimal chromosomal abnormalities

We next characterized the selected iSTOP hiPSC clones for stem cell pluripotency, chromosomal abnormalities, and neuron differentiation capability. Immunofluorescence staining of stem cell pluripotency markers (OCT4, SOX2, TRA-1-60) of the engineered hiPSC lines for 6 selected LoF alleles all confirmed their pluripotency (Figures 4A, S4A). To further evaluate the pluripotency of all the selected hiPSC LoF clones at the genomic and molecular level, we carried out RNA-seq for each hiPSC clone and used CellNet to quantify how closely the engineered hiPSC populations transcriptionally resembled human embryonic stem cells (ESC) compared to other non-ESC somatic cells (Cahan et al., 2014). All hiPSC clones exhibited high stemness scores (0.93∼0.97) and no traces of other somatic cell types (Figures 4B, S4B, S4C). With the same RNA-seq data, we also confirmed the absence of large chromosomal abnormalities using eSNP-Karyotyping (Weissbein et al., 2016; Zhang et al., 2023) (Figures 4C, S5). Finally, we confirmed that all the selected iSTOP hiPSC lines (n=6) could be successfully induced into neurons (MAP2+/Syn) after Ngn2-transduction (Figure 4D).

**Figure 4.**
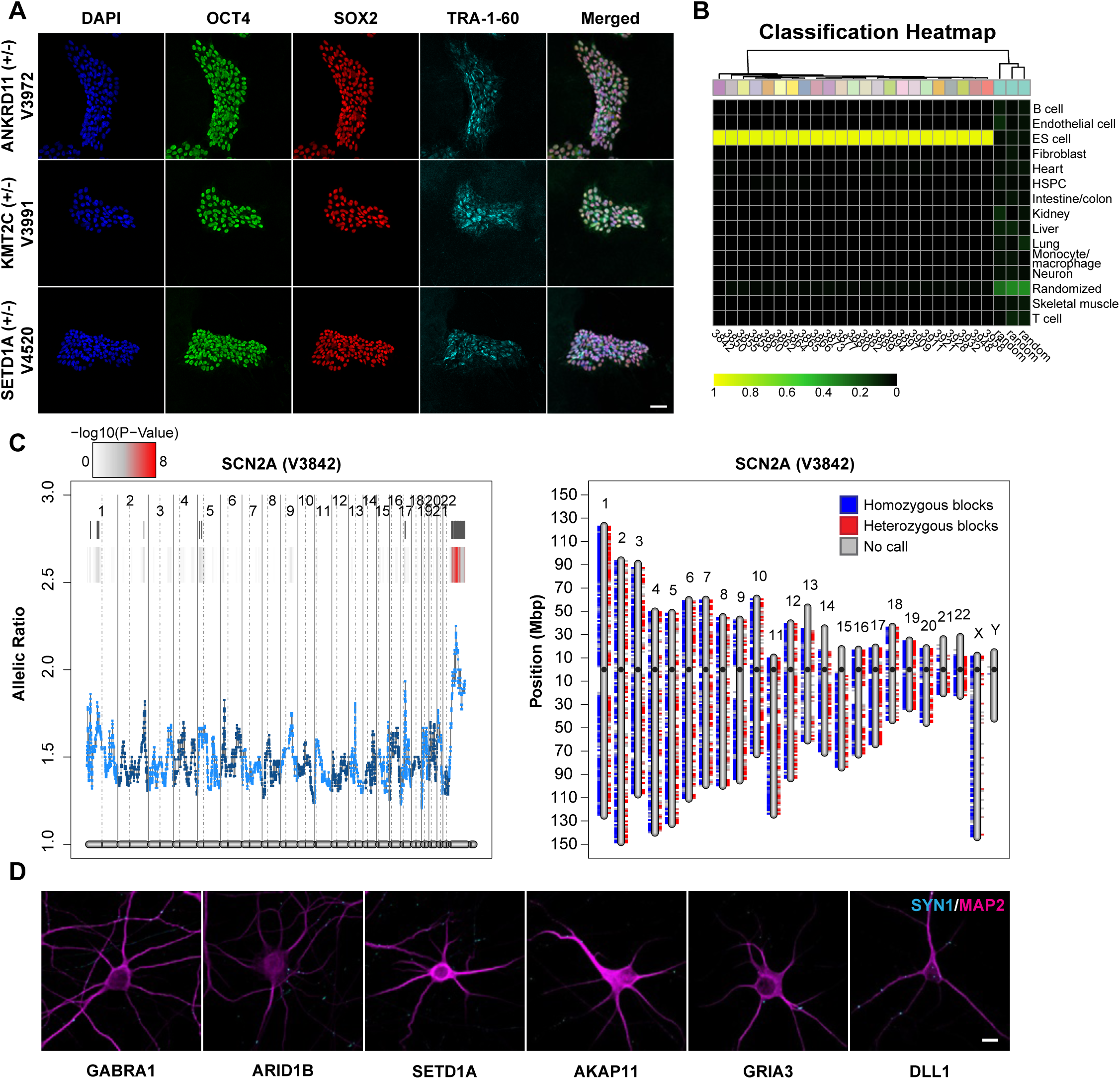
Characterization of isogenic base-edited hiPSC lines carrying iSTOP LoF alleles. (A) The iSTOP mutant lines were stained positive for pluripotent stem cell markers (OCT4, SOX2, TRA-1-60). Scale bar: 50µm. (B) CellNet analysis of RNA-seq data of hiPSC lines confirmed their pluripotency. Pluripotency scores showed transcriptional similarity of the edited iSTOP LoF hiPSC lines to ESC or other non-ESC cell types. Only one batch of hiPSC lines are shown, data of the other two batches are in Figure S4. (C) e-Karyotyping showed no large chromosomal abnormalities. Example for one hiPSC line is shown. (D) Some selected iSTOP LoF hiPSC lines were successfully differentiated into excitatory neurons (Syn+/MAP2+). Scale bar: 10µm. In (A) and (C), the gene name for the LoF allele and the cell line number (starting with “V”) were listed.

### Most iSTOP hiPSC lines show the expected mRNA or protein reduction with the confirmation of SHANK3 LoF phenotype in Ngn2-induced neurons

Because we have employed the iSTOP approach to introduce premature stop codons that are predicted to cause NMD (Table S1), we first tested whether we could observe the expected expression reduction for each NPD gene in the engineered hiPSC lines using RNA-seq data. Compared to the unedited cell line, the iSTOP lines for about 12 genes showed partial or near-complete expression knockdown (KD) as expected for NMD (Figures 5A, S6). Strongest expression KD was observed for hiPSC lines homozygous for *SCN2A, CHD8* and *CACNA1G* iSTOP LoF alleles, exhibiting a 70-90% expression reduction. The lack of the expected NMD for some genes may be due to possible cell type-specific NMD regulation (Huang et al., 2011), incomplete mRNA degradation, or inaccurate NMD prediction in sgRNA design. Our qPCR further confirmed the incomplete mRNA degradation, or even increased in mRNA production (e.g., *HCN4*, *SP4*) for genes that did not show the expected NMD in RNA-seq (Figure 5B).

**Figure 5.**
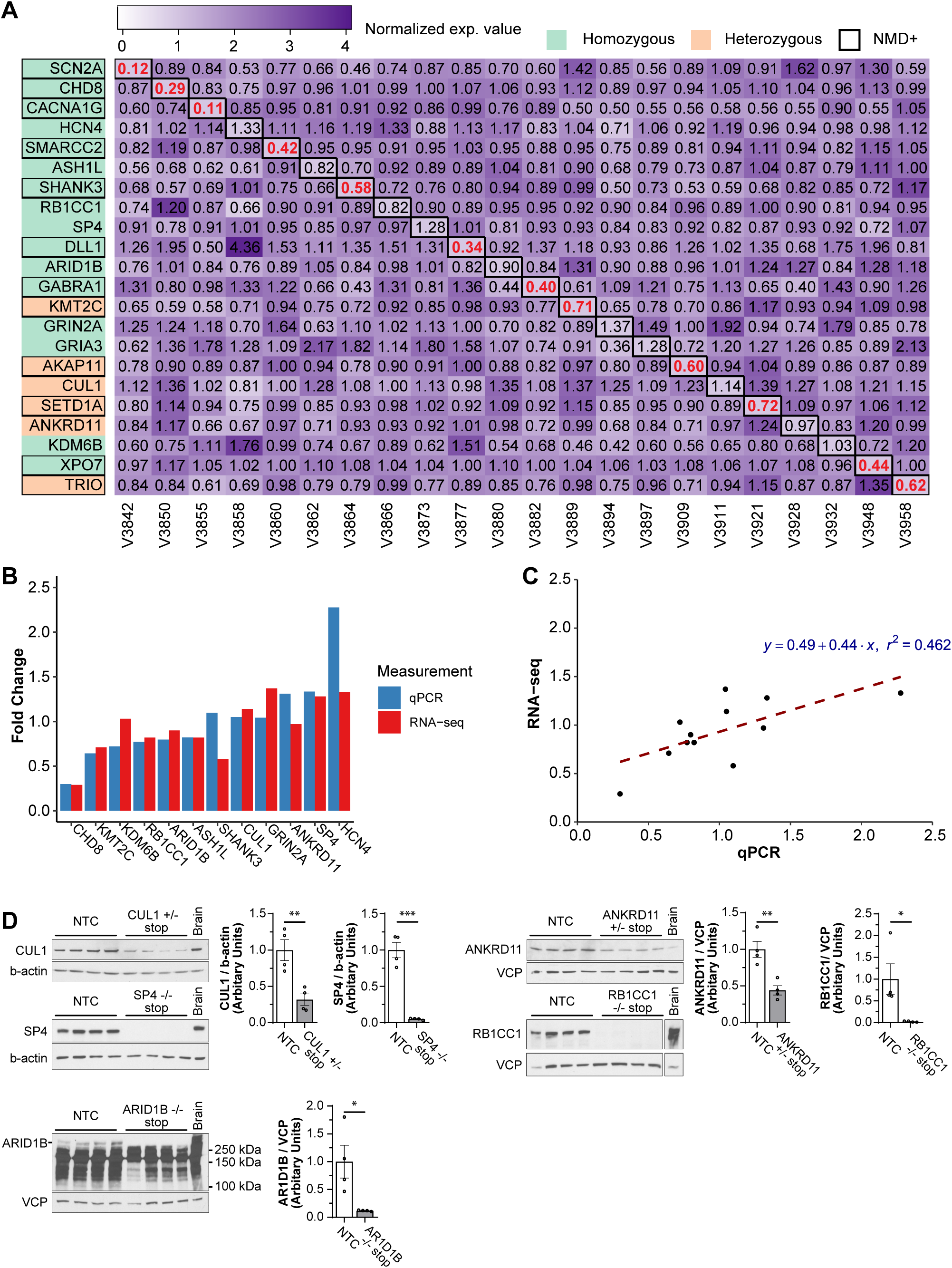
Characterization of NMD and loss-of-function for iSTOP LoF hiPSC lines. (A) Heatmap of expression fold changes of each mutant line (vs. NTC) using normalized RNA-seq expression value (CPM, count per million). The values in the diagonal boxes show the fold-change of a specific line with LoF mutation. The fold-change values in red fonts indicate those showing NMD. (B) qPCR confirmation of the expression fold change in RNA-seq (vs. NTC). (C) Strong correlation of expression fold changes (vs. NTC) between RNA-seq and qPCR data. (D) Western blots showed the expected reduction of protein abundance for the LoF alleles of 5 selected genes that did not exhibit NMD in (A) and (B). NTC=non-transfected control. Mouse brain protein extracts were used as a positive control for each blot. Note that non-specific signal below 250 kDa, expected sizes for ARID1B, were apparent in the blots, only putative signal of ARID1B was quantified. Two-tailed Student’s *t*-test with unequal variance was used for comparison between unedited and LoF lines, n = 4 biological replicates (different cell cultures). * *P* <0.05, ** *P* < 0.01, *** *P* < 0.001.

Regardless of any detectable NMD from RNA-seq or qPCR, we expected those premature stop-codons at the first half of a target gene would result in protein truncations (i.e., the loss of full-length proteins). To confirm this hypothesis, we performed Western blotting for 5 selected genes that did not show the expected mRNA NMD (*CUL1, ANKRD11, SP4, RB1CC1, ARID1B*) (Figures 5A, 5B) using cell lysates of their respective iSTOP hiPSC lines (*CUL1* +/-, *ANKRD11* +/-, *SP4* -/-, *RB1CC1* -/-, *ARID1B* -/-) (Figure 5C). We found that compared to the unedited hiPSC line (NTC), all 5 LoF lines showed the expected protein reduction based on their genotype, with heterozygous LoF lines showing ∼50% decrease of the intact proteins (CUL1, ANKRD11) while homozygous LoF lines exhibiting near complete KD (SP4, ARID1B, RB1CC1) (Figure 5C). This result strongly suggests that most LoF alleles engineered by our iSTOP base editing approach effectively led to the expected gene expression KD or complete abolishment of the gene expression, i.e., a LoF effect.

To further corroborate the LoF effect by the introduced iSTOP mutation, we assessed whether we could replicate the previously reported morphological phenotypes in *SHANK3*-deficient human neurons (Yi et al., 2016). We derived excitatory (Ex) and inhibitory (Inh) induced neurons by ectopic expression of Ngn2 or Ascl1/Dlx2 transcription factors (Halikere et al., 2020; McGowan et al., 2018; Yang et al., 2017; Zhang et al., 2013) (see Methods) from the hiPSC lines that carry homozygous iSTOP LoF allele of *SHANK3*. We also included an edited hiPSC line that carried the iSTOP LoF allele of *CUL1*, a strong SZ risk gene (Singh *et al*., 2022b) that only had heterozygous clones despite its high editing efficiency (Figure 3B). With the co-cultures of Ex- and Inh-neurons, we assayed the neurite outgrowth, branches and synaptic puncta density of the tdTomato-labeled Ex neurons using high content imaging (HCI). Compared to the Ex-neurons from the isogenic control hiPSC line, the *SHANK3* iSTOP LoF line showed ∼2/3 reduction of neurite outgrowth and branches but no significant change of synaptic puncta (Synapsin1+) density (Figures 6, S7), which are consistent with the reported cellular phenotypes of S*HANK3*-haploinsufficiency in human neurons (Yi *et al*., 2016). For the SZ risk gene *CUL1*, we observed a significant reduction of Ex-neuronal neurite outgrowth and branches by ∼60% as well as a reduced synaptic puncta (Synapsin1+) density by ∼40% (Figures 6, S7).

**Figure 6.**
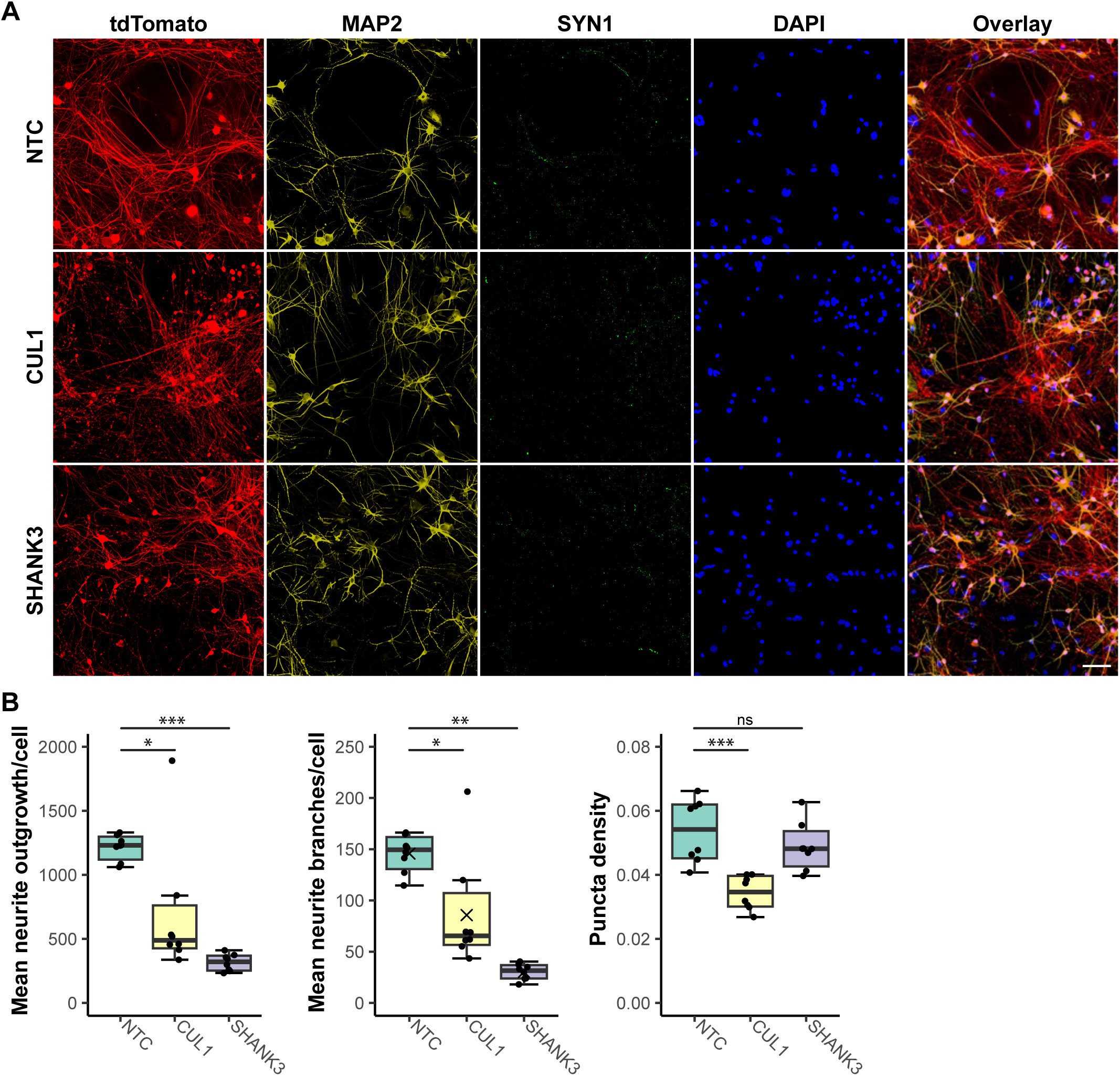
High-content imaging of excitatory and inhibitory neurons co-cultured with mouse glia for iSTOP LoF hiPSC lines. (A) Representative images of neural cultures with hiPSC-differentiated excitatory neurons labeled by tdTomato (red). Neurons were stained for MAP2 and SYN1, tdTomato and DAPI. TdTomato and DAPI staining were used for quantifying neurite growth/branches; tdTomato, MAP2 and SYN1 staining were used for synaptic puncta analysis. Scale bar: 20µm. (B) Summarized imaging result for neurite outgrowth and branches and synaptic puncta (left to right) for SHANK3 (-/-) and CUL1 (+/-) LOF alleles on donor hiPSC line CW2017. Each data point represents measurement of a cell culture replicate in a well of a 96-well plate. N=8 replicates from a single experiment. Two-tailed Student’s t-test with unequal variance was used for testing for differences between NTC and a LoF allele group. * *P* <0.05, ** *P* < 0.01, *** *P* < 0.001, ns = not significant.

Taken together, our observed NMD of mRNA, protein expression reduction, and neural phenotypic changes in the assayed iSTOP hiPSC lines suggest that most iSTOP hiPSC lines for NPD genes are expected to show LoF effects.

## Discussion

Although CRISPR editing of individual NPD risk genes/variants in hiPSC has been widely used in the past decade (De Los Angeles *et al*., 2021; Duan, 2023; Michael Deans and Brennand, 2021; Muhtaseb and Duan, 2022; Wang *et al*., 2020; Wen *et al*., 2016; Zhang *et al*., 2023; Zhang et al., 2020), a scaled and efficient pipeline for clonal LoF mutagenesis in hiPSC has not been established. Our reported CBE iSTOP editing workflow benefited from the improved gene editing efficiency of the CBEmax_enrich system, the precision of iSTOP mutagenesis, the streamlined RNA-seq-based assays for pluripotency, eSNP-Karyotyping, and iSTOP-mediated NMD and/or LoF. These factors simplified the workflow and made it more amenable for automation to increase throughput while keeping the pipeline cost-effective. Furthermore, barcode tracking, semi-automation, and well-controlled editing (i.e., by including an editing control in every batch of 23 target genes with one NTC control line) ensured unbiased and robust assays of cellular and molecular phenotypes of LoF alleles. Moreover, because our pipeline only involved transient transfection of hiPSC, the engineered iSTOP hiPSC lines were genome-integration-free, as opposed to CRISPR pooled screening that often entails hiPSC genome-integration with exogenous virus fragments that may confound downstream phenotypic assay readouts. The derived iSTOP hiPSC lines carrying LoF alleles for the current list of 22 (out of 23) edited genes on 3 donor hiPSC lines, and many more engineered LoF hiPSC lines to be generated, will be an invaluable sharable resource for the NPD genetics research community.

The scaled CBE iSTOP editing pipeline enabled us to systematically test CBE editing efficiency in different hiPSC lines for many genes. We observed high editing efficiency for most genes that was highly reproducible across all three hiPSC lines, suggesting CBE iSTOP performance was not hiPSC line-specific, but rather mainly determined by gene-specific sgRNA design. As expected, genes with high editing efficiency tended to have more clones homozygous for iSTOP LoF alleles, and all four genes with relatively low editing efficiency (<10%) gave only heterozygous clones. However, and interestingly, for three strong SZ risk genes (*SETD1A*, *CUL1, TRIO*) identified by SCHEMA we only obtained heterozygous LoF hiPSC clones despite high editing efficiency, suggesting the likely deleterious effect of LoF on stem cell survival. Indeed, possible lethal effects of LoF of the three genes is supported by the existing body of literature: *Setd1a*, encoding a histone methyltransferase, was found to be required for embryonic and neural stem cell survival (Bledau et al., 2014), and only heterozygous *SETD1A*-haploinsufficiency hiPSC lines (Chong et al., 2022; Wang et al., 2022; West et al., 2019) or mouse models (Mukai et al., 2019; Nagahama et al., 2020) have been reported for functional characterization; *CUL1* has E3 ubiquitin-protein ligase activity and homozygous deletion of *Cul1* in mice causes arrest in early embryogenesis (Wang et al., 1999); *TRIO* functions as a guanosine diphosphate (GDP) to guanosine triphosphate (GTP) exchange factor and is necessary for cell migration and growth (Deinhardt *et al*., 2011; Seipel *et al*., 1999). Therefore, regardless of the CRISPR editing tool (CRISPR/Cas9 or CBE), a proportion of the selected NPD genes are likely to only have heterozygous LoF alleles in hiPSC, for which a later-stage inducible editing system would be required to assay the phenotypic effects of homozygous LoF. On the other hand, it is noteworthy that highly penetrant patient-specific PTVs in these NPD genes are all heterozygous. Thus, the obtained heterozygous LoF hiPSC clones can still be valuable for ascertaining more disease-relevant cellular phenotypes.

Our clonal LoF mutagenesis pipeline leverages the precise control of the CBE iSTOP editing approach to introduce a premature stop-codon that is predicted to cause NMD of mRNAs or lead to a protein truncation. However, as shown in our systematic editing of the 22 NPD genes, only 12 showed partial or nearly complete NMD in hiPSC. Other than possible inaccuracy of NMD prediction, the lack of the expected NMD for some genes are likely due to: 1) RNA-seq may have still detected partially degraded mRNAs; 2) an intricate feedback network maintains both RNA surveillance and the homeostasis of normal gene expression in mammalian cells (Huang *et al*., 2011); 3) some cells do escape NMD either by translational readthrough at the premature stop codon or by a failure of mRNA degradation after successful translation termination (Sato and Singer, 2021). However, it is noteworthy that, as confirmed by our Western blotting analysis, even without detectable NMD most iSTOP mutations, if not all, likely led to “LoF” by yielding a truncated protein too short to be detected by Western blot. Although a truncated protein may arguably have normal function or even “gain” of function, the sgRNA in our iSTOP design often targets the protein N-terminal, thus more likely to show LoF. Though translational readthrough occasionally produces a full length of protein, such translational readthrough is rare (Sato and Singer, 2021). It is reassuring that even for genes that do not show NMD (e.g., *CUL1*) or even increased mRNA level (likely due to negative feedback regulation, e.g., *SP4*) in hiPSC, we have observed the expected protein expression KD or complete protein expression knockout as demonstrated by Western blotting and moreover, a robust reduction of neurite growth/branches or synaptic puncta density in the SHANK3 LoF neurons, suggesting most iSTOP mutant alleles are likely to show LoF in neurons.

Despite overall high efficiency, our CBE iSTOP editing efficiency remained low for a few genes (5 out of 22). Further improvement may include optimization of sgRNA design and the use of multiple iSTOP sgRNAs. Among other limitations, our iSTOP design relies on NMD or protein truncation to achieve LoF, where the extent of “LoF” may not be as complete as obtained from complete KO using CRISPR/Cas9. However, a complete gene KO may require the use of multiple gRNAs targeting multiple genomic regions, which poses a challenge for clonal LoF mutagenesis on a large scale and, more importantly, may cause large chromosomal arrangements. Lastly, although many genes can be edited in parallel, the throughput of our clonal iSTOP LoF mutagenesis pipeline remains relatively low, which may be improved by pooled CRISPR screening in combination with the rapidly evolving spatial transcriptomics and phenotyping to simultaneously ascertain LoF allelic identity and assay cellular phenotypes at single neuron resolution. Albeit limitations, our scaled and efficient clonal LoF mutagenesis pipeline showed robust performance in generating easily sharable individual hiPSC lines carrying LoF alleles for a large number of NPD genes. In addition to engineering LoF alleles, the pipeline can be easily adopted for precise SNP editing (C to T changes by CBE, and A to G changes by ABE) in hiPSC for functional characterization of either coding or noncoding disease risk variants. Moreover, our modified PAM-less CBEmax (and ABE) system further expands the repertoire of the CRISPR gene editing toolbox, empowering hiPSC as a promising cellular model for understanding the disease biology of NPD and other complex genetic disorders.

## Experimental Procedures

### Resource availability Lead contact

Further information and requests for resources and reagents should be directed to and will be fulfilled by the lead contact, Jubao Duan

### Materials availability

The plasmids generated in this report will be made available on request but with a completed Materials Transfer Agreement. The iPSC lines generated in this study will be made available as part of SSPsyGene consortium to fulfill the NIMH (National Institute of Mental Health) material/data sharing commitment; however, due to the current lack of and external centralized repository for its distribution and our need to maintain the stock, we are glad to share the cell lines in the form of collaboration according to NIMH instructions.

### Data and code availability

All the reported data and the original code for using RNA-seq data to test for pluripotency will be made available upon request.

### hiPSC lines and cell culture

The use of CW20107 (from California’s Stem Cell Agency - CIRM) and KOLF2.2J (from The Jackson Laboratory) was part of the SSPsyGene consortium agreement on the common cell lines. The other 4 hiPSC lines (CD14, CD19, 8565612726, 8129019249) were specific to the MiNND project and were from Duan lab (Shi *et al*., 2009; Zhang *et al*., 2023; Zhang *et al*., 2020). Only 3 (CW20107, KOLF2.2J and CD14) of the 6 cell lines were used in the current report. Detailed cell line information is described in Table S2. The hiPSCs were maintained in mTeSRPlus (StemCell #100-0276) with primocin (Invitrogen #ant-pm-1) on tissue culture plates coated with matrigel (Fisher Scientific #08-774-552) or geltrex (Fisher Scientific #A1413202) throughout the mutagenesis process. The Institutional Review Board (IRB) of NorthShore University HealthSystem approved study.

### Gene selection and iSTOP base editing design

The reported 23 genes were part of the ∼250 NPD genes selected by SSPsyGene consortium (<u>sspsygene.ucsc.edu</u>). The various gene selection criteria included a strong association with NPD, mainly SZ and ASD, with highly penetrant rare coding risk variants. For creating LoF alleles, we used an improved DNA base editing system (iSTOP C to T editor) to efficiently introduce premature stop codons (i.e., nonsense mutations) that resemble the highly penetrant NPD risk variants expected to cause NMD and/or protein truncation. For designing iSTOP sgRNAs for each gene, we first retrieved the best pre-computed sgSTOP-RNA for each selected NPD gene using the iSTOP webtool (Billon *et al*., 2017), requiring >50% NMD rate and in >50% transcript isoforms. NMD prediction was determined based on whether the targeted base was 55 nucleotides upstream of the final exon-exon junction (Billon *et al*., 2017; Popp and Maquat, 2016). Whenever possible, the sgSTOP location was placed in the first half of the gene to ensure the resultant protein truncation (likely to be LoF) even without causing NMD. For cloning the designed sgRNAs, pDT-sgRNA (Addgene# 138271) vector was selected as gRNA carrier, and the Gibson assembly approach was used to clone sgRNAs into the vector.

### iSTOP base editing pipeline

The cell culture, DNA base editing/sgRNA transfection, and cell sorting for LoF mutagenesis were performed in batches, each containing 23 genes and a non-transfected control (NTC) on a 24-well plate format. The base editor system (pEF-AncBE4max, pEF-BFP, and pDT-sgRNA) contains a reporter gene that makes cells that have undergone C to T editing turn from blue to green . LipofectamineSTEM was used for cell transfection. After cell transfection for 72 hrs, single hiPSC from the post-transfection culture were sorted into 96-well plates with one cell per well using a BD FACSAria Fusion Flow Cytometer in the presence of CEPT cocktail (1:10,000 chroman 1, emricasan, and transISRIB; 1:1,000 polyamine supplement) (Tristan *et al*., 2023). The sorted single cells were cultured on a 96-well plate with media changes every other day for 10-14 days until colonies appeared with an appropriate size to pick for Sanger sequencing genotyping. About 8-12 colonies were picked for each editing condition. Sanger sequencing on 3730xl DNA Analyzer was completed to confirm that the appropriate base was changed at the desired location to create a stop codon. SeqScape v2.5 was used for automatic DNA sequencing analysis and genotype calling. Up to 4 colonies with confirmed homozygous editing or heterozygous editing (if there were no homozygous colonies) and good morphology were expanded for RNA isolation and hiPSC banking.

### hiPSC characterization (quality control and LoF confirmation)

The selected LoF mutant lines for banking were characterized for stem cell pluripotency by both immunofluorescence staining for pluripotency markers (for a few cell lines) and transcriptomic characterization by using CellNet analysis of RNA-seq data of each isogenic hiPSC line (Cahan *et al*., 2014). For characterizing possible hiPSC chromosomal abnormality, as we previously described (Zhang *et al*., 2023; Zhang *et al*., 2020), we used e-Karyotyping (Weissbein *et al*., 2016). For characterizing the effects of iSTOP mutation on NMD, we analyzed RNA-seq data of each hiPSC line to quantify the expression value of each targeted gene and compare to its expression value in the NTC line. For some selected genes, we also confirmed NMD and the loss of intact proteins in LoF lines by qPCR and Western blotting, respectively.

### Neuron differentiation from hiPSC

We used the two commonly used methods for differentiating iPSC lines into excitatory and inhibitory neurons: Ngn2 + rtTA for excitatory neuron differentiation (Zhang *et al*., 2013), and Ascl1 + Dlx2 + rtTA for inhibitory neuron differentiation (Yang *et al*., 2017). In brief, Lentiviral vectors were generated by transfecting HEK293T cells with lentivirus packaging plasmids (pMDLg/pRRE, VsVG and pRSV-REV) with the desired vectors as previously described (Pang et al., 2011). hiPSCs were dissociated with Accutase, cells were counted and 2e5 cells were plated per well in 6-well plates coated with Matrigel. A mixture of virus was added to the cell media before plating: i) Ngn2 + rtTA was added for excitatory neuron differentiation (Zhang *et al*., 2013), ii) Ascl1 + Dlx2 + rtTA was added for inhibitory neuron differentiation (Yang *et al*., 2017). Excitatory neurons were also transduced with a lentivirus with a plasmid expressing TdTomato on day 4 to distinguish them from inhibitory neurons. On day 5 induced neurons were dissociated with Accutase and counted, and then co-cultured with mouse glia. Half the media was changed every 5 days. On day 35, cells were washed and fixed with 4% PFA for 30 min. Cells were left in PBS 0.02% sodium azide until staining.

### Neuron morphological characterization

For immunofluorescence staining, hiPSC-derived neurons were fixed in a 96 well optical bottom plate with a polymer base (Fisher Scientific: 12-566-70) permeabilized. The neurons were stained with primary antibodies, mouse anti-Synapsin 1 (1:500), goat anti-tdTomato (1μg/ml), and chicken anti-MAP2 (1:5000), and incubated with appropriate secondary antibodies. For Image acquisition, the neurons were imaged using Molecular Devices (San Jose, CA) ImageXpress Micro Confocal High-Content Imaging System at both 20× and 40×. The mean number of neurite branches per cell and the mean length of neurite outgrowth per cell were analyzed with the built-in Neurite Outgrowth Application Module within the MetaXPress 6 software, version 6.7.2.290. For assaying excitatory synapse density, we used an in-house generated custom synaptic assay module with MetaXPress 6 software. The puncta density was generated by the number or total area of puncta within the colocalized MAP2 and tdTomato staining divided by the area of MAP2+ & tdTomato+ signal within the neurites.

## Acknowledgments

Data was generated as part of the SSPsyGene Consortium, supported by RM1MH133065 awarded to Z.P.P., J.D., and J.M. We thank SSpyGene consortium members for selecting NPD genes and providing iPSC lines CW20107 (from CIRM) and KOLF2.2 (from the Jackson Lab). We thank Molecular Genetics of SZ (MGS) investigators for collecting samples that were used to derive MGS hiPSC lines. We also thank A.R.S. for collecting samples for deriving iPSC bank of neurodegenerative disorders (iBOND) and Genomic Health Initiative (GHI). Funding: Funding was provided by NIH grants R01AA023797 and R01MH125528 (to Z.P.P.), R01MH106575, R01MH116281, and R01AG063175 (to J.D.); and RM1MH133065 (to Z.P.P., J.D., and J.M.).

## Author Contributions

H.Z, L.P, and A.M. designed the mutagenesis pipeline, performed the main experiments and wrote the manuscript; S.D.D.L.G performed the neuron differentiation and characterization, wrote the manuscript; S.Z. analyzed the RNA-seq data and wrote the manuscript; P.G. and M.E.A. performed neuron differentiation and characterization; D.S. and G.T. performed the neuron imaging analyses and wrote the manuscript; A.D. assisted with RNA-seq data analysis; W.W. and B.J. assisted with hiPSC derivation and culture; R.P. assisted with the project coordination; R.H., C.P., J.M., and A.R.S. assisted with project design, result interpretation, and edited the manuscript; Z.P.P. and J.D. conceived and supervised the project, and wrote the manuscript.

Supplemental information (Supplementary Figures, Tables and Experimental Procedures) is provided.

