## Supplementary Information for "Scaled and Efficient Derivation of Loss of Function Alleles in Risk Genes for Neurodevelopmental and Psychiatric Disorders in Human iPSC"

#### Supplementary Figures

Figure S1.

**A**

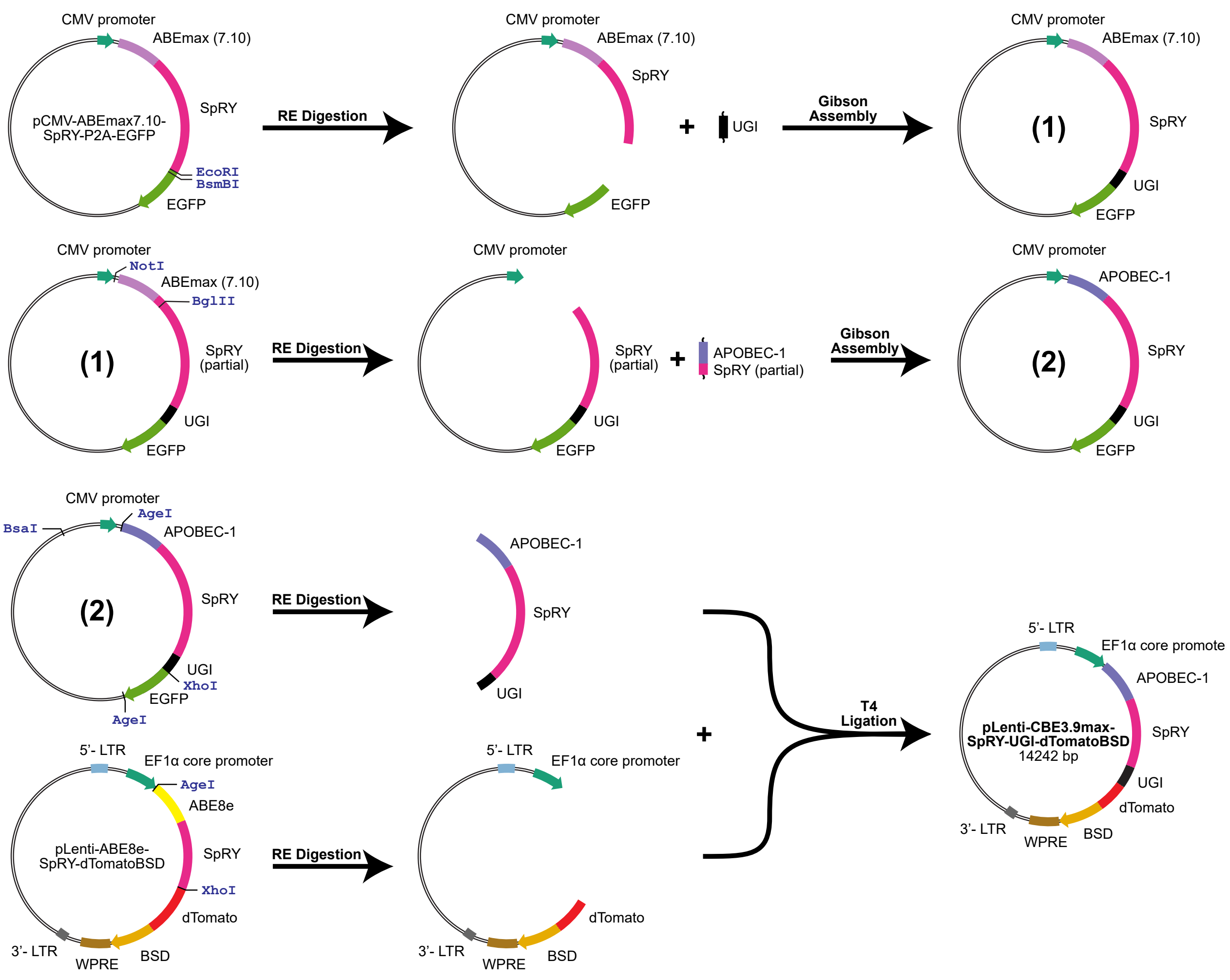
CMV promoter
ABEmax (7.10)
pCMV-ABEmax7.10-SpRY-P2A-EGFP
SpRY
EcoRI
BsmBI
EGFP
RE Digestion
UGI
Gibson Assembly
(1)
NotI
BglII
SpRY (partial)
APOBEC-1
(2)
BsaI
AgeI
XhoI
5'- LTR
EF1α core promoter
ABE8e
pLenti-ABE8e-SpRY-dTomatoBSD
dTomato
3'- LTR
WPRE
BSD
T4 Ligation
pLenti-CBE3.9max-SpRY-UGI-dTomatoBSD
14242 bp

**B**

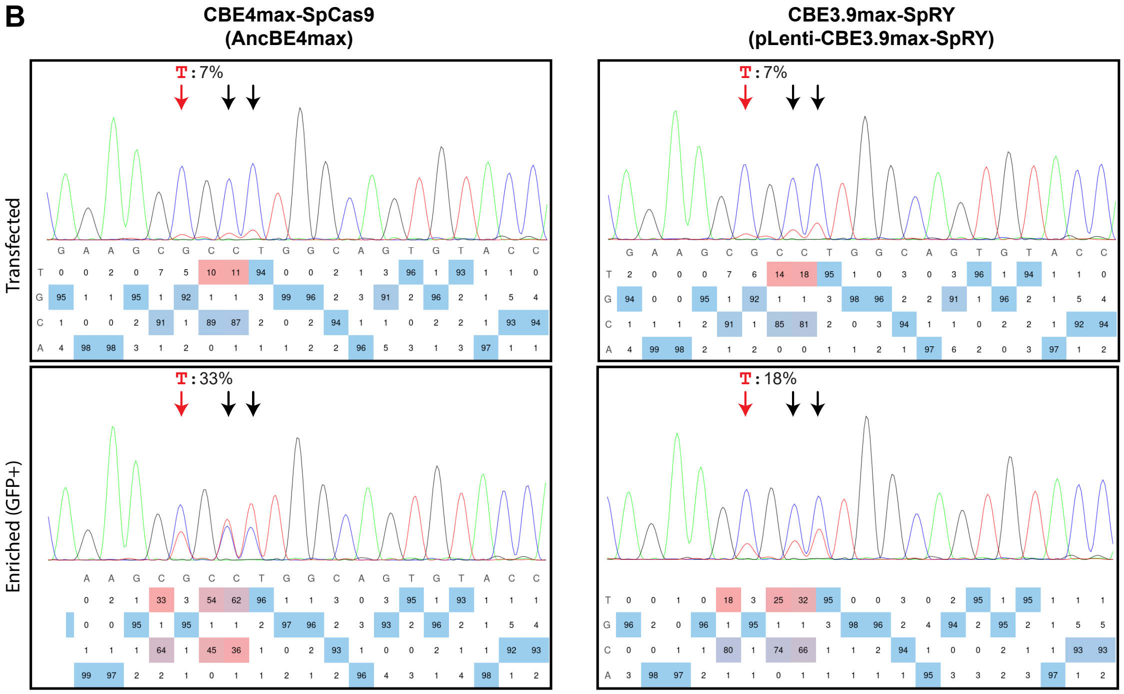
CBE4max-SpCas9
(AncBE4max)
CBE3.9max-SpRY
(pLenti-CBE3.9max-SpRY)
Transfected
Enriched (GFP+)
T: 7%
T: 7%
T: 33%
T: 18%

**Figure S1**. An improved PAM-less CBE base editor *pLenti-CBE3.9max-SpRY-UGI-dTomatoBSD* (associated with Figure 1). (A) Schematics of the plasmid DNA vector construction for a PAM-less CBE base editor *pLenti-CBE3.9max-SpRY-UGI-dTomatoBSD.* (B) C to T editing efficiency in hiPSC at a target site before and after editing enrichment (reporter GFP+ cells) using regular PAM-based *CBE4max-SpCas9* and *PAM-less CBE3.9max-SpRY*. By-standing editing events were highlighted with black arrow; however, for iSTOP mutations, such by-sanding editing are in protein codon(s) proceeding to the  iSTOP site thus would not affect the LoF effect of an iSTOP mutation.

Figure S2

A

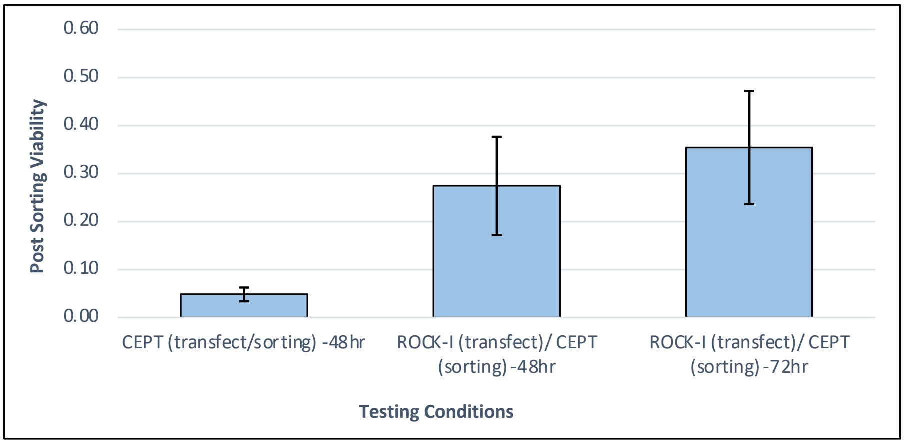


B

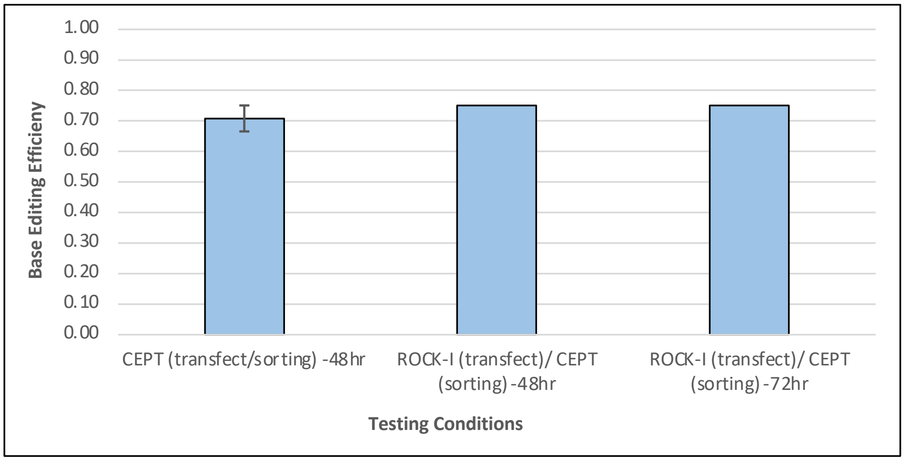


C

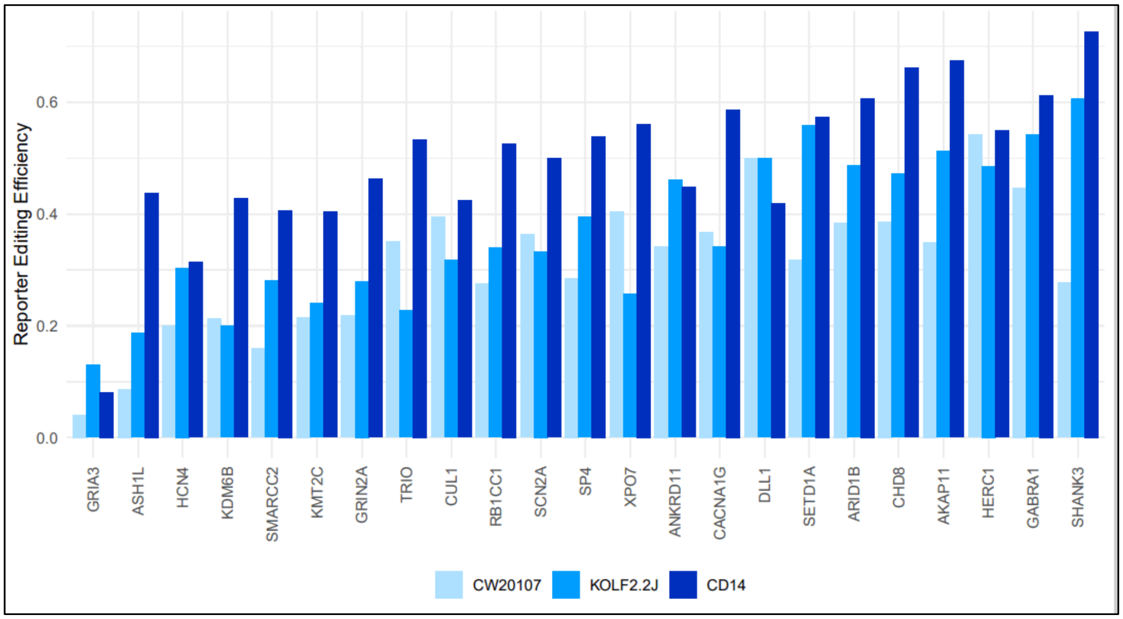

**Figure S2.** Reporter gene editing efficiency and optimization for increasing post-sorting single hiPSC clonal survivability (associated with Figures 2 and 3). (A) Post-sorting single hiPSC clonal survivability (A) and C to T base editing efficiency (B) (assayed by Sanger sequencing of the LoF mutation site) under different transfection/cell sorting conditions. (C) Reporter gene editing efficiency for all the target genes in three hiPSC lines (CW20107, KOLF2.2J, and CD14).

Figure S3.

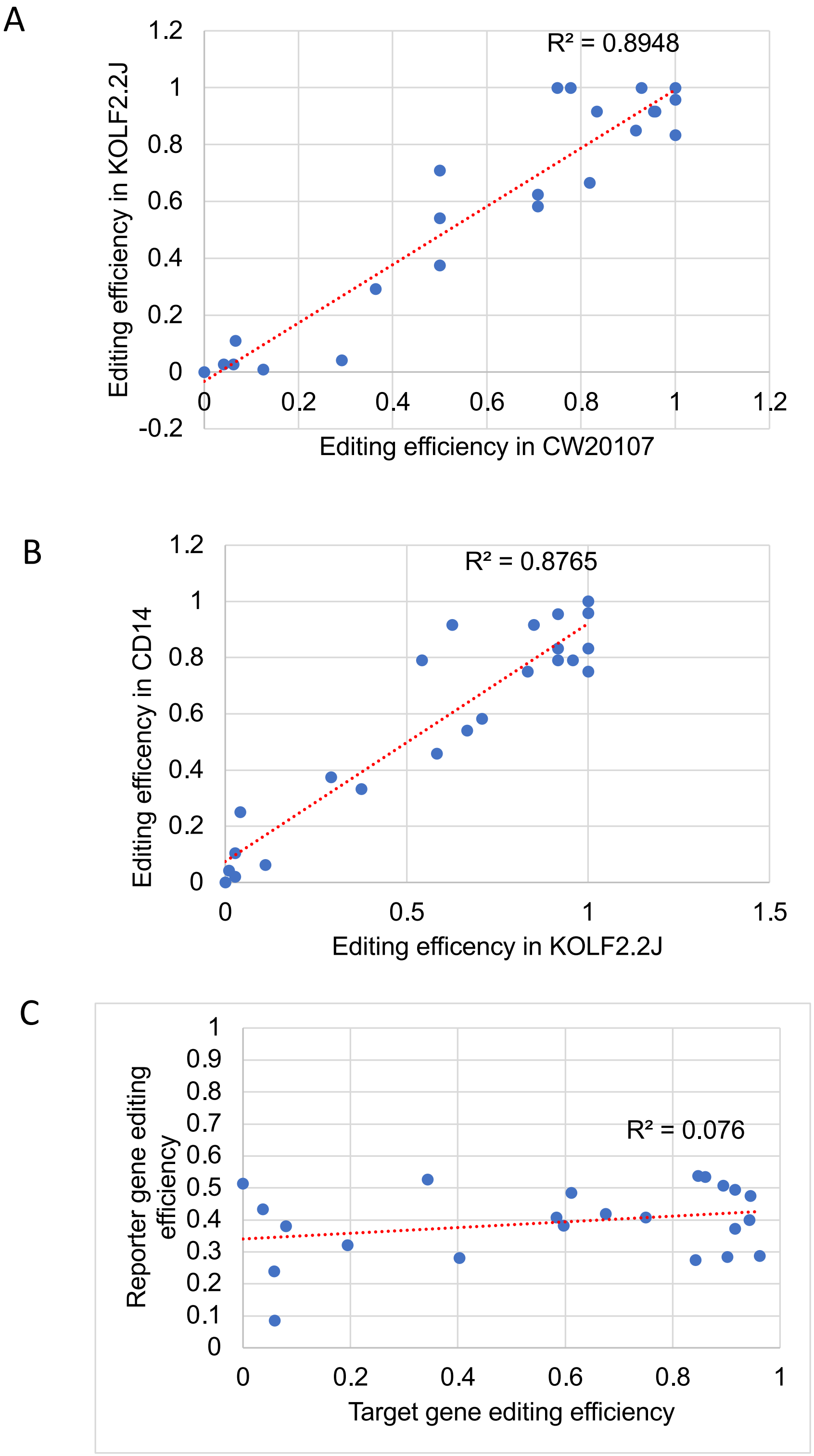
A
R² = 0.8948
Editing efficiency in KOLF2.2J
Editing efficiency in CW20107
1.2
1
0.8
0.6
0.4
0.2
0
-0.2
0
0.2
0.4
0.6
0.8
1
1.2
B
R² = 0.8765
Editing efficency in CD14
Editing efficency in KOLF2.2J
1.2
1
0.8
0.6
0.4
0.2
0
0
0.5
1
1.5
C
R² = 0.076
Reporter gene editing efficiency
Target gene editing efficiency
1
0.9
0.8
0.7
0.6
0.5
0.4
0.3
0.2
0.1
0
0
0.2
0.4
0.6
0.8
1

**Figure S3.** Pearson correlation of editing efficiency between different hiPSC lines (associated with Figure 3). (A) and (B) Strong correlations of target gene editing efficiency across cell lines. (C) Weak correlation between target gene editing efficiency and reporter gene editing efficiency. Target gene editing efficiency was calculated by the genotypes of each single hiPSC clone as confirmed by Sanger sequencing. Report gene editing efficiency in (C) was calculated as the proportion of GFP+ cells (reporter gene edited cells) vs. BFP+ cells (cells transfected with sgRNAs) in FACS.

Figure S4.

A

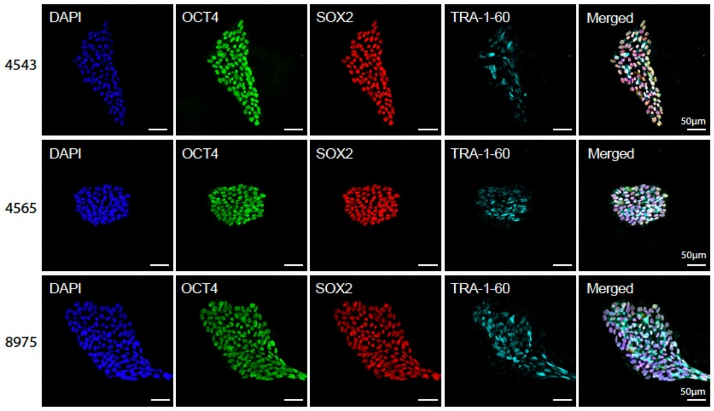


B

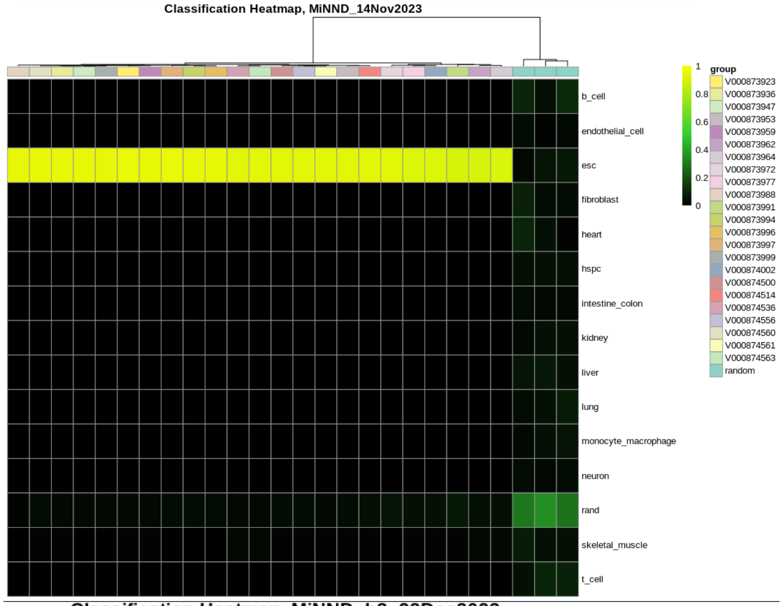


C

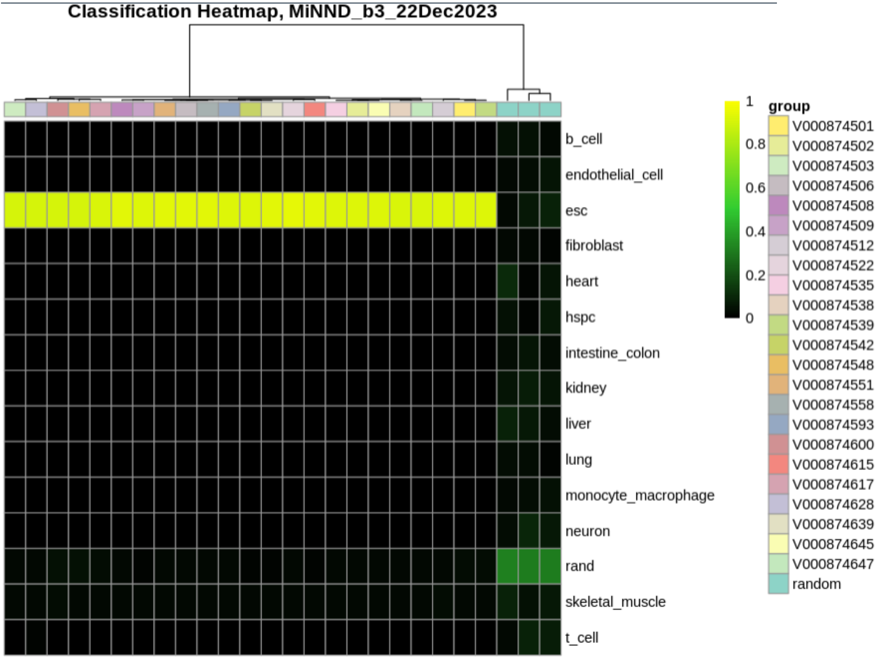

**Figure S4**. Stem cell pluripotency characterization of iSTOP LoF hiPSC lines (associated with Figure 4). (A) The iSTOP mutant lines were stained positive for pluripotent stem cell markers (OCT4, SOX2, TRA-1-60). Scale bar: 50μm. (B) CellNet analysis of RNA-seq data of hiPSC lines confirmed their pluripotency. Pluripotency scores showed the transcriptional similarity of the edited iSTOP LoF hiPSC lines to ESC or other non-ESC cell types. Two batches of edited hiPSC lines are shown.

### Figure S5.

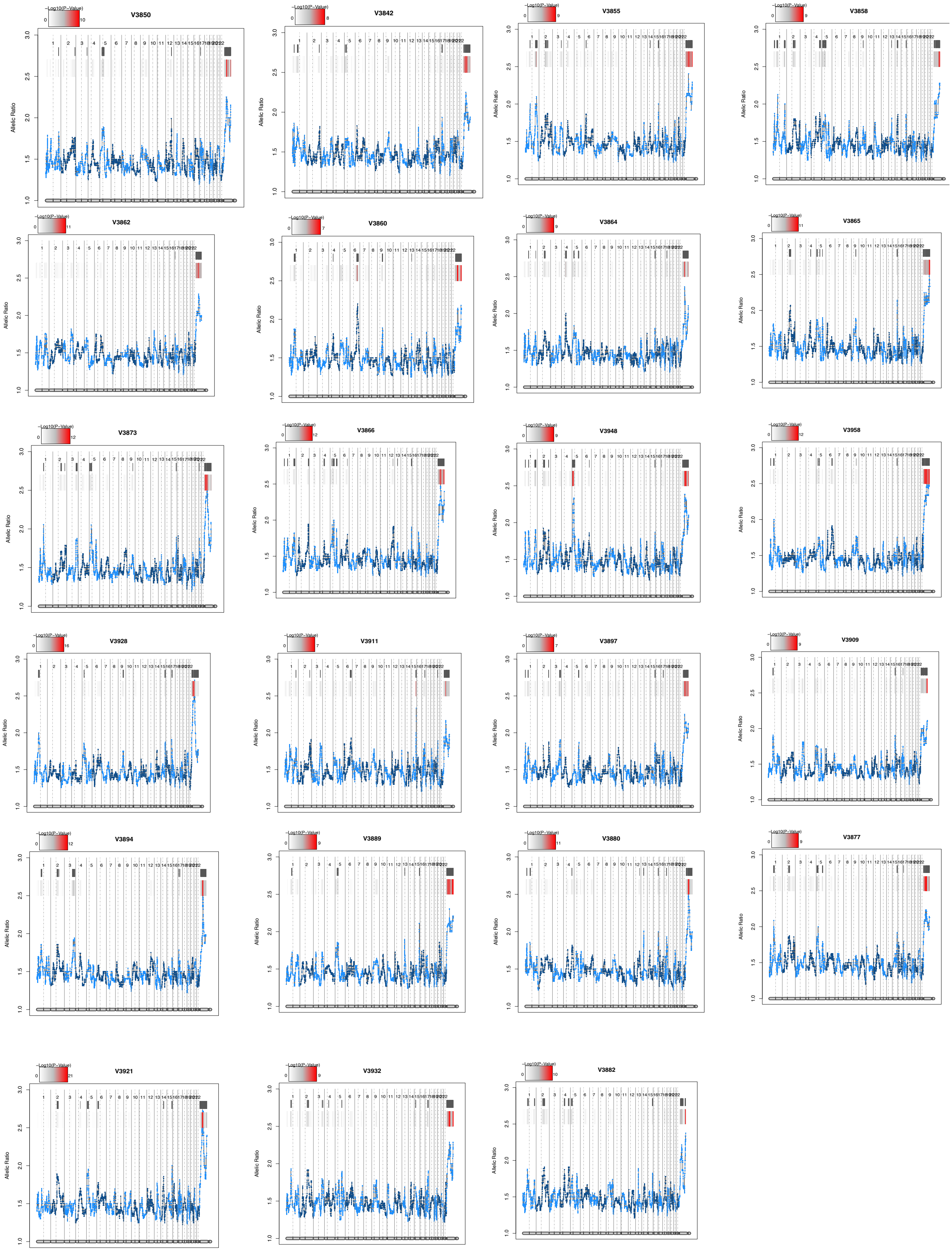
V3850
V3842
V3855
V3858
V3862
V3860
V3864
V3865
V3873
V3866
V3948
V3958
V3928
V3911
V3897
V3909
V3894
V3889
V3880
V3877
V3921
V3932
V3882
-Log10(P-Value)
Allelic Ratio

**Figure S5.** RNA-seq based eSNP-Karyotyping (only batch 1 edited lines are shown) (associated with Figure 4). Each panel showed the moving average of SNP intensity (RNA-seq reads) of the two alleles of heterozygous SNPs.

Figure S6.

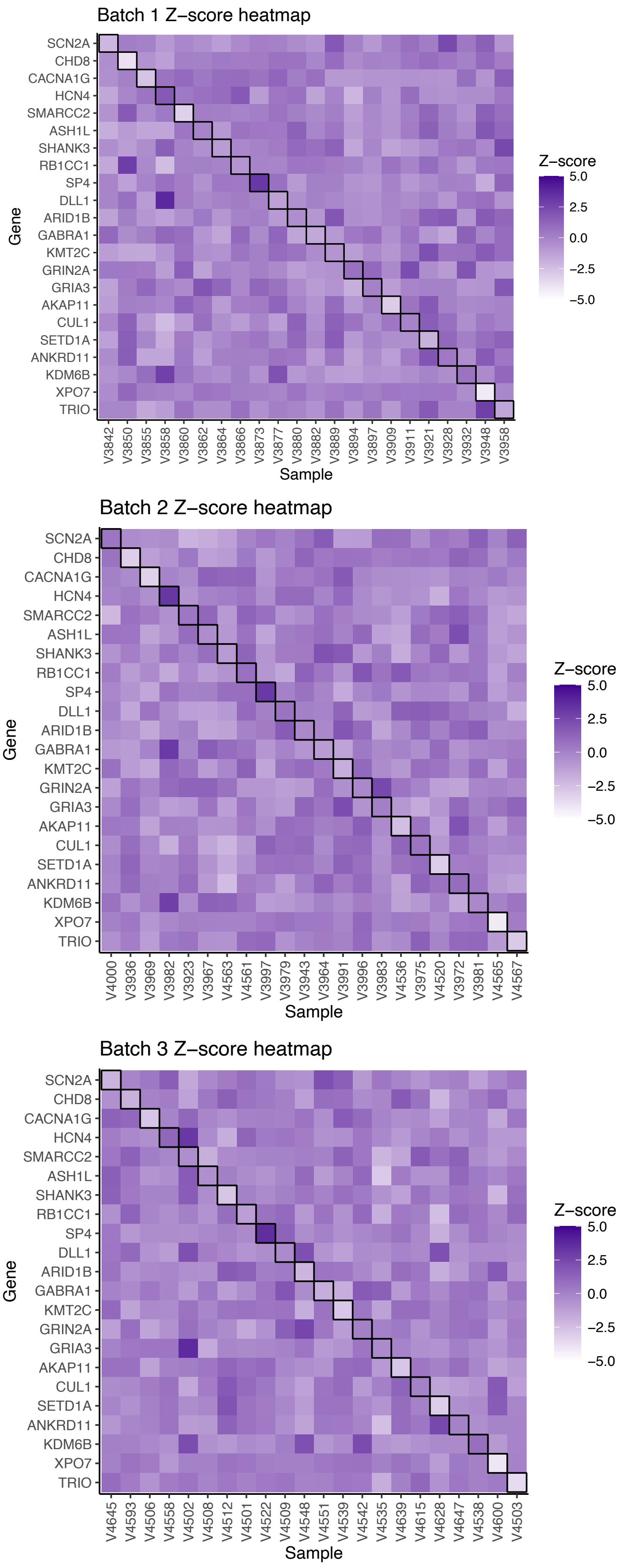
Batch 1 Z−score heatmap
Gene
SCN2A
CHD8
CACNA1G
HCN4
SMARCC2
ASH1L
SHANK3
RB1CC1
SP4
DLL1
ARID1B
GABRA1
KMT2C
GRIN2A
GRIA3
AKAP11
CUL1
SETD1A
ANKRD11
KDM6B
XPO7
TRIO
V3842 V3850 V3855 V3858 V3860 V3862 V3864 V3866 V3873 V3877 V3880 V3882 V3889 V3894 V3897 V3909 V3911 V3921 V3928 V3932 V3948 V3958
Sample
Z−score
5.0
2.5
0.0
−2.5
−5.0
Batch 2 Z−score heatmap
Gene
SCN2A
CHD8
CACNA1G
HCN4
SMARCC2
ASH1L
SHANK3
RB1CC1
SP4
DLL1
ARID1B
GABRA1
KMT2C
GRIN2A
GRIA3
AKAP11
CUL1
SETD1A
ANKRD11
KDM6B
XPO7
TRIO
V4000 V3936 V3969 V3982 V3923 V3967 V4563 V4561 V3997 V3979 V3943 V3964 V3991 V3996 V3983 V4536 V3975 V4520 V3972 V3981 V4565 V4567
Sample
Z−score
5.0
2.5
0.0
−2.5
−5.0
Batch 3 Z−score heatmap
Gene
SCN2A
CHD8
CACNA1G
HCN4
SMARCC2
ASH1L
SHANK3
RB1CC1
SP4
DLL1
ARID1B
GABRA1
KMT2C
GRIN2A
GRIA3
AKAP11
CUL1
SETD1A
ANKRD11
KDM6B
XPO7
TRIO
V4645 V4593 V4506 V4558 V4502 V4508 V4512 V4501 V4522 V4509 V4548 V4551 V4539 V4542 V4535 V4639 V4615 V4628 V4647 V4538 V4600 V4503
Sample
Z−score
5.0
2.5
0.0
−2.5
−5.0

**Figure S6**. RNA-seq-based estimation of the expected NMD for each iSTOP LoF allele derived from all three donor hiPSC lines (associated with Figure 5). Shown are heatmaps of Z-scored expression values (counts per million reads, CPM) of each gene in each iSTOP LoF hiPSC line. Lined boxes indicate the edited hiPSC line that is expected to show NMD for a specific target gene.

Figure S7.

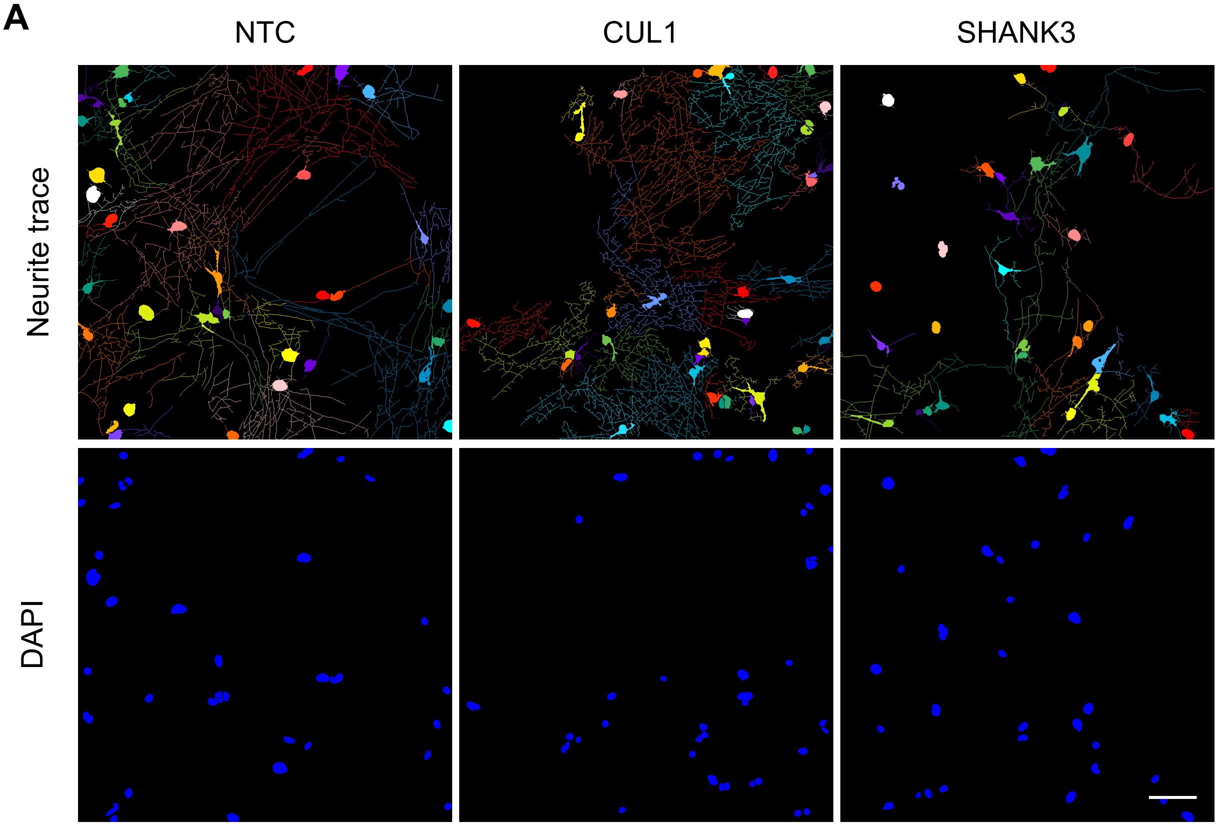


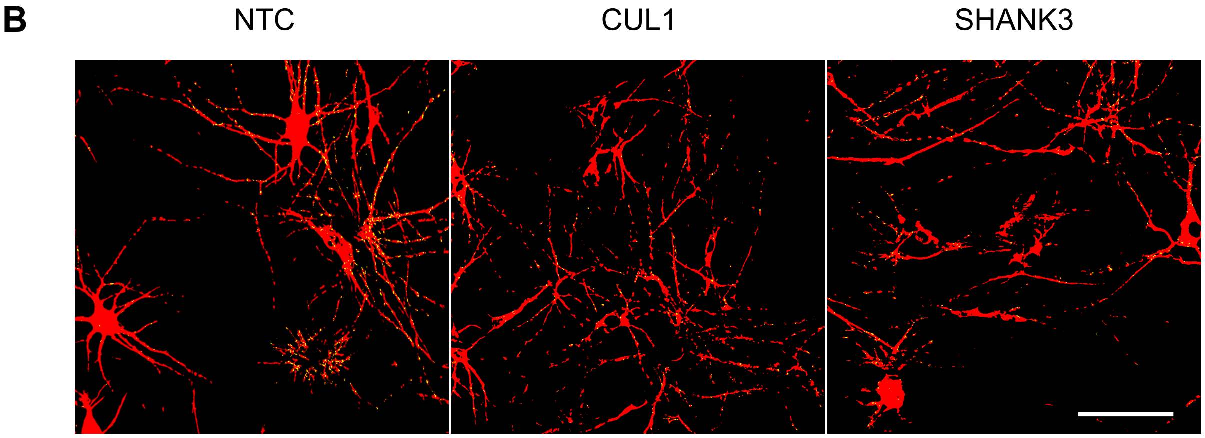

**Figure S7**. Cell segmentation and binary masks in morphometric analysis using high content imaging in neuron/glia co-culture (associated with Figure 6). (A) Cell segmentation for assaying neurite outgrowth and branches of LoF alleles of SHANK3 (-/-) and CUL1 (+/-) using the build-in neurite outgrowth module on the ImageXpress system. Single neurons were highlighted in rainbow color. (B) Binary masks for assaying puncta density of LoF alleles of SHANK3 (-/-) and CUL1 (+/-) using a customized synaptic module. MAP2+ and tdTomato+ mask was highlighted in red, and SYN1+ mask was highlighted in yellow. Scale bar in all panels: 20μm.

#### Supplementary Tables

**Table S1**. Sequences and other genomics information for iSTOP sgRNAs

**Table S2**. Cell lines for generating LoF alleles (EA=European ancestry, AA=African ancestry)

**Table S3**. Genotypes at the iSTOP site for each LoF hiPSC line (NA=not available; no editing)

**Table S4**. Oligos and primers used for sgRNA cloning, sequencing confirmation, and qPCR assays

**Supplemental Experimental procedures**

**1. The source hiPSC lines**

CW20107 was purchased from California's Stem Cell Agency (CIRM). KOLF2.2J was purchased from The Jackson Laboratory. Two other hiPSC lines (CD14 and CD19) of European ancestry were derived by National Institute of Mental Health (NIMH) Stem Cell Center (also known as RUCDR-Rutgers University Cell & DNA repository) from donors of the Molecular Genetics of Schizophrenia (MGS) cohort (Shi et al., 2009), and have been characterized for pluripotency, absence of large chromosomal abnormality, and neural differentiation in our previous studies (Zhang et al., 2023; Zhang et al., 2020). The two hiPSC lines of African ancestry (8565612726 and 8129019249) were derived as part of the iBOND (induced pluripotent stem cell bank of neurodegenerative disorders) cohort in house at NorthShore University HealthSystem, and have undergone similar characterization to that of MGS hiPSC lines (Zhang et al., 2023; Zhang et al., 2020). More detailed information on donor cell lines can be found in Table S2. The Institutional Review Board (IRB) of NorthShore University HealthSystem approved study.

**2. Chemicals and reagents**

The chemicals, media, and reagents used in cell culture, PCR, Sanger sequencing, and other main experiments include: BbsI-HF (NEB: R3539S), NEBuilder® HiFi DNA Assembly reagent (NEB: E2621S), mTeSR Plus (StemCell: 100-0276), Primocin (InvivoGen: ant-pm-1), Matrigel (FisherScientific: 08-774-552), Geltrex (Fisher Scientific: A1413202), DPBS (no calcium, no magnesium) (Fisher Scientific: 14-190-144), ReLeSR (StemCell: 100-0483), ROCK-Inhibitor (Tocris: 1254), mFreSR (Stem Cell: 05855), Accutase (StemCell: 07920), Trypan Blue Stain (FisherScientific: T10282), Gibco Opti-MEM I Reduced Serum Medium (Fisher Scientific: 31-985-062), Lipofectamine Stem Transfection Reagent (Fisher Scientific: STEM00003), Chroman 1 (R&D Systems: 7163/10), Emricasan (R&D Systems: 7310/5), transISRIB (R&D Systems: 5284/10), Polyamine Supplement (R&D Systems: 7739/1), QuickExtract DNA Extraction Solution 1.0 (Fisher Scientific: NC9904870), PCRx Enhancer System (Fisher Scientific: 11-495-017), Deoxynucleoside Triphosphate Set (Sigma Aldrich: 3622614001), Shrimp Alkaline Phosphatase and buffer (Fisher Scientific: 783905000UN), E.coli exonuclease I (Fisher Scientific: 70073X5000UN), BigDye Terminator v3.1 Cycle Sequencing Kit (Fisher Scientific: 4337455), HiDi Formamide (Fisher Scientific: 4311320), RNeasy Plus Mini Kit (Qiagen: 74134), QIAshredder (Qiagen: 79654), DMSO (Sigma Aldrich: D2650-100ML), Fetal Bovine Serum (Fisher Scientific: A3160501), Mineral Oil (Sigma Aldrich: M5310), 16% Formaldehyde (Fisher Scientific: PI28908), DAPI (Fisher Scientific: EN62248). The antibodies used in immunofluorescence staining include: Synapsin I antibody (Synaptic Systems: 106-011), goat anti-tdTomato antibody (Fisher Scientific: 50-167-1115), anti-MAP2 antibody (Sigma Aldrich: AB5543), Donkey Anti-Mouse 488 (Fisher Scientific: A21202), Donkey Anti-Goat IgG (H+L) Cross Adsorbed Secondary Antibody, Alexa Fluor 568 (Fisher Scientific: A11057), Donkey Anti-Chicken IgY (H+L) Highly Cross Adsorbed

Secondary Antibody, Alexa Fluor 647 (Fisher Scientfic: A78952). The main lab supplies include: 96-well non skirted PCR plate (DotScientific: 650-PCR), Corning™ Internally Threaded Cryogenic Vials 2 ML (FisherScientific: 03-374-21), 96-well plates (Corning: 353072), 24-well plates (ThermoFisher: 142475), 6-well plates (StemCell: 38016).

**3. iSTOP gRNA design and cloning**

We first retrieved the best pre-computed sgSTOP-RNA for each selected NPD gene using iSTOP webtool (Billon et al., 2017), requiring >50% NMD rate and in >50% transcript isoforms. NMD prediction was determined based on whether the targeted base was 55 nucleotides upstream of the final exon-exon junction (Billon *et al.*, 2017; Popp and Maquat, 2016). Whenever possible, the sgSTOP location was placed to the first half of the gene to ensure the resultant protein truncation (likely to be LoF) even without causing NMD.  Finally, to minimize any possible off-target editing, all the selected sgSTOPs are free of any predicted off-target site via aligning to the human genome, allowing up to two mismatches in the first eight bases of the guide sequence (Billon *et al.*, 2017; Popp and Maquat, 2016). A total of 23 genes were designed for sgSTOP-RNAs in the current study (See Table S1). For cloning the designed sgRNAs, pDT-sgRNA (Addgene# 138271) vector was selected as gRNA carrier. The vector was digested with BbsI-HF. After gel purification, a single strand oligo with prefix, gRNA of interest and post fix was introduced into the vector backbone through Gibson assembly using NEBuilder® HiFi DNA Assembly reagent. After cloning, mini prep plasmid was sequenced using M13 Rev primer. M13 Rev: 5' – caggaaacagctatgac – 3'. Example of a single strand oligo: 5' – atatcttgtggaaaggacgaaacaccgXXXXXXXXXXXXXXXXXXXXgttttagagctagaaatagcaagtta – 3'. After genotyping, correct clones were expanded and transfection grade plasmid was prepared using endo-free plasmid kit (QIAGEN: 12362).

**4. Lenti-CBEmax-SpRY construction**

Cloning was started from pCMV-T7-ABEmax(7.10)-SpRY-P2A-EGFP (Addgene#140003). Vector was digested with EcoRI-HF and BsmBI, then UGI unit was introduced into gel purified vector through Gibson assembly to generate the first intermediate plasmid pCMV-ABEmax(7.10)-SpRY-UGI-P2A-EGFP. The plasmid DNAs of pCMV-ABEmax(7.10)-SpRY-UGI-P2A-EGFP were subsequently digested with NotI-HF and BglII to remove adenine base editor unit. Then the cytosine base editor unit APOBEC-1 was introduced into the digested backbone through Gibson assembly, which further gave the second intermediate plasmid pCMV-CBE3.9max-SpRY-UGI-P2A-EGFP. The pCMV-CBE3.9max-SpRY-UGI-P2A-EGFP was then digested with AgeI-HF, XhoI, BsaI-HFv2, and the part of CBE3.9max-SpRY-UGI insert was purified; at the same time, the pLenti-ABE8e-SpRY-dTomatoBSD plasmid was digested with AgeI-HF and XhoI to obtain the backbone. CBE3.9max-SpRY-UGI insert was then ligated into the pLenti-ABE8e-SpRY-backbone though T4 DNA ligation, giving rise to the final plasmid pLenti-CBE3.9max-SpRY-UGI-dTomatoBSD. All the intermediate plasmid sequences were validated with Sanger sequence before proceeding next steps.

**5. HEK293 culture and transfection**

HEK293T cells were purchased from ATCC (Cat# CRL-3216) and maintained in DMEM with 10% FBS following vendor's instructions. For transfection and editing efficiency evaluation, 90% confluent 293T culture was dissociated with accutase at 37°C for 5min. About $2\times10^5$ cells were replated into one well on 12-w tissue culture plate. 24hr post replating, 2μg of selected CBE plasmid, 1μg of pEF-BFP (Addgene# 138272) plasmid and 1μg of pDT-sgRNA plasmid carrying selected gRNA were transfected using Fugene HD (Promega# E2311) reagent with 1:3 DNA:Reagent ratio following vendor's instructions. 48hr post transfection, BFP+/GFP+ and BFP+/GFP- cells were sorted through BD Aria Fusion Flow Cytometer and replated. 120hr post transfection, replated cells were collected after accutase dissociation and 30-75μl QuickExtract DNA Extraction Solution (FisherSci # NC9904870) was added to the cell pellet for DNA extraction on thermocycler. Extracted DNA was subsequently amplified for Sanger sequencing genotyping to evaluate editing efficiency at loci of interest.

**6. hiPSC culture and transfection**

hiPSCs were maintained in mTeSRPlus (StemCell #100-0276) with primocin (Invivogen #ant-pm-1) on tissue culture plates coated with matrigel (Fisher Scientific #08-774-552) or geltrex (Fisher Scientific #A1413202) throughout the mutagenesis process. Medium was changed every other day and colonies were passaged every 4-6 days when cells reached 70% confluence. For DNA transfection, cells were plated at a density of $1.2\sim1.5\times10^5$ per well on a 24-well plate (ThermoFisher # 142475) in mTeSR Plus with 5μM ROCK-Inhibitor (Tocris #1254). The next day, antibiotics-free mTeSR Plus with 5μM ROCK-Inhibitor was changed on the plate after ensuring appropriate cell density (60-70% confluence) and survival. Shortly after, each well was transfected with 750ng pEF-AncBE4max (addgene #138270), 300ng pEF-BFP, and 300ng pDT-sgRNA containing the variant-specific gRNA using LipofectamineSTEM with 1:2.5 DNA:reagent ratio. In total, 23 different pDT-sgRNAs were used (one per well) and the 24th well was used as a negative control. Media was refreshed with regular mTeSR Plus containing primocin at 24hr and 48hr post transfection. At 72hr post transfection, cells were prepped for single cell sorting.

**7. Single hiPSC sorting and clonal culture**

Single hiPSC from the post-transfection culture above were sorted into 96 well plates with one cell per well using a BD FACSAria Fusion Flow Cytometer. All sorting procedures were done using mTeSR Plus with

CEPT cocktail (1:10,000 chroman 1, emricasan, and transISRIB; 1:1,000 polyamine supplement) (Tristan et al., 2023). To prepare for FACS, cells were dissociated into single cells using accutase (StemCell: 07920) for 7min at 37°C. Cells were transferred to 15ml tubes with 1ml mTeSR plus to inactivate the accutase and centrifuged at 300×g for 3min. The resulting cell pellets were resuspended in 700μl media and filtered twice using 5ml corning round bottom tubes with blue strainer cap (Fisher Scientific: 0877123). Samples were placed on ice immediately to minimize clogging. Samples were processed and analyzed using the BD FACSAria Fusion Flow Cytometer, gating for BFP+/GFP+ cells, one single cell per well on a 96 well plate for each condition; for NTC, BFP+/GFP- cells were sorted. Following sorting, plates were centrifuged at 300×g for 1 min to aid in cell attachment, then returned to the incubator. After sorting, the cells were not disturbed for 24hr. 48hr post sorting, 50μl mTeSR plus was added to the each well. 72hr post sorting, 50μl media refreshment for each well. 96hr post posting, 120μl media refreshment for each well. 144hr post sorting, aspirated 120μl and added 100μl media for each well. Afterwards, 100μl media refreshment every other day (10-14 days) until colonies appeared with an appropriate size to pick for Sanger sequencing genotyping. All media refreshments were performed using Integra's MINI 96 electronic pipette.

##### 8. PCR and Sanger sequencing for LoF genotype confirmation

Once colonies reached an appropriate size and had stem cell-like morphology, 8-12 colonies were picked for each edited condition. DNA was extracted from the picked colonies using Quick Extract DNA Extraction Solution (Fisher Scientific NC9904870). Following PCR to amplify the DNAs for each LoF gene, Sanger sequencing was completed to confirm that the appropriate base was changed at the desired location to create a stop codon. The sequencing was performed on a 3730xl DNA Analyzer and the sequencing data were imported to SeqScape v2.5 for automatic analysis and genotype calling. Up to 4 colonies with confirmed homozygous editing or heterozygous editing (if there were no homozygous colonies) and good morphology were expanded for RNA isolation and cell cryopreservation.

##### 9. RNA isolation for RNA sequencing (RNA-seq)

Based on sequencing results, two selected clones were passaged from one well on 96-well plates to one well on 6-well plates. Once reaching 70% confluency, cells were expanded a second time from one well to two wells per clone, one for cryopreservation and one for RNA extraction/RNA-seq. For RNA isolation, cells were lysed using 800μl Buffer RLT Plus (QIAGEN 1053393). Cell lysates in buffer RLT were stored at -80°C until ready to be isolated using the QIAGEN RNeasy Plus Mini Kit (QIAGEN 74134) following vendor's instructions. Purified RNAs were sent to Novogene for RNA-seq.

**10. RNA-seq data processing**

Bulk RNA-seq were performed by external vendor Novogene and were provided in 2×150 bp paired-end format of 25-30 M reads per sample. Briefly, raw FASTQ files were aligned to the human GRCh38.p13 genome by STAR 2.7.0 and subsequently counted by the in-build function of STAR at the gene level using the GTF file of GENCODE v30 with parameters --quantMode Genecounts --alignSoftClipAtRefenceEnds No --outFilterScoreMinOverLread 0.30 --outFilterMatchNminOverLread 0.30. Gene counts from each of the samples were collected by a customized script and collated into a single count matrix. Genes that had 0 counts in all samples were removed prior to analysis.

**11. CellNet analysis for pluripotency**

The RNA-seq data of each edited iSTOP hiPSC line were used for pluripotency evaluation by using the R package CellNet (Cahan et al., 2014). Briefly, the gene × sample count matrix generated in the previous step was loaded by EdgeR and a new count matrix contained log-transformed, library size-normalized CPM (counts-per-million) value was generated by calcNormFactors(), estimateDisp() and cpm(). Subsequently, the script constructed a random forest classifier using the in-built model from the CellNet Package. Finally, the likelihood of each sample-cell type pair (in scores) was evaluated by passing the log-transformed gene × sample count matrix through the classifier and plotting the results in hierarchically clustered heatmaps.

**12. Using RNA-seq data for e-SNP Karyotyping**

As part of the high throughput LoF mutagenesis pipeline for evaluating possible hiPSC chromosomal abnormality due to editing or hiPSC clonal growth, we opted to use e-SNP Karyotyping rather than the classical G-band karyotyping. e-SNP Karyotyping detects any potential chromosomal aberrations, including duplications, loss of heterozygosity, and meiotic recombination. As we previously described (Zhang *et al.*, 2023; Zhang *et al.*, 2020), we used e-Karyotyping R package developed by the Benven lab (github.io/BenvenLab/eSNPKaryotyping) (Weissbein et al., 2016) with customization to our current environment settings. The same BAM file set generated from bulk RNA-seq using STAR 2.7.0 was used. The original code was optimized to reflect the updated R (4.3.1), GATK (4.2.6.1), and dbSNP 154 was used in our study.

**13. Immunofluorescence staining for hiPSCs**

hiPSCs were dissociated with Accutase (innovative cell technologies AT-104) and seeded into Matrigel (Cornig 354234) coated round glass coverslips in a 24 well plate and kept in mTESR+ media (Stem cell technology 100-0275). Cells were kept until they formed medium sized colonies. Cells were washed twice with 1× PBS and fixed with 4% PFA for 30 min. Samples were incubated with blocking buffer 4% BSA (A3803 Sigma), 1% Goat serum (Thermo Fischer 16210072), 0.2% triton X-100 (BP151 Fisher BioReagents) in PBS for 1hr at room temperature (RT). Primary antibodies were incubated for 1hr at RT. Samples were washed 3 times with PBS 0.2% Triton X-100 and secondary antibodies were incubated for 1hr at RT making sure samples were protected from light. Samples were washed 3 times with PBS 0.02% Triton X-100 and rinsed with MiliQ water before mounting with Fluoroshield with DAPI (F6057 Sigma) and placed on a glass slide. The following antibodies were used: rabbit anti-Sox2 (Millipore AB5603), mouse IgG anti-Oct4 (Millipore MAB4401), mouse IgM anti-Tra-1-60 (Millipore MAB4360), goat anti-rabbit Alexa Fluor 546 (Invitrogen A11035), goat anti-mouse IgG Alexa Fluor 488 (Invitrogen A11001), goat anti-mouse IgM Alexa Fluor 647 (Invitrogen A21238).

**14. Western blot**

hiPSC cells were grown on 6 well plates as stated above. When the cells reached ~80% confluency, they were washed twice with DPBS and lysed with 100µl RIPA buffer (50 mM Tris-HCl pH 7.5, 150 mM NaCl, 1% NP-40, 0.5% sodium deoxycholate, 0.1% SDS) supplemented with 0.5mM DTT, 1mM PMSF and 1× Protease inhibitor Cocktail (Sigma P8340) using a cell scraper. The lysate was transferred to a tube incubated on ice for 10 min and centrifuged at 14000×g for 10 min at 4°C. The supernatant was transferred to a new tube and protein concentration was determined using a BCA protein assay (Thermoscientific 23225) measuring absorbance at 562nm in a plate reader (SpectraMax i3, Molecular Devices). For the SDS-PAGE, protein samples were prepared with 15µg of protein with 2× Laemmli sample buffer (161-0737 BIORAD) and heated for 5 min at 95°C. Samples were resolved using 7.5% or 10% acrylamide gels (4561023DC, 4561033DC BIORAD) and transferred into a 0.45µm nitrocellulose membrane (1620115 BIORAD). Membranes were blocked for 1hr at RT using 5% non-fat milk dissolved in TBS-T (20 mM Tris-HCl pH 7.5, 150 mM NaCl, 0.1% Tween 20). Primary antibodies were diluted in 5% non-fat milk (732-291-1940 LabScientific) or 5% BSA (A3803 Sigma-Adrich) in TBS-T and incubated overnight at 4°C. Membranes were washed 3 times with TBS-T and incubated with an HRP-conjugated secondary antibody diluted in 5% non-fat milk or BSA in TBS-T for 1hr at RT. Membranes were washed 3 times with TBS-T and protein was visualized using the Clarity Western ECL Substrate (Biorad 1705060) and autoradiography films (XAR ALF 1318, LabScientific). The following antibodies were used: BAF250b/ARID1B (1:3000 Cell Signaling 92964S), beta-Actin (1:20000 Sigma-Aldrich A544), CUL1 (1:2000 Santa Cruz SC-17775), FIP200/RB1CC1 (1:2000 Cell Signaling 12436S, SP4 (1:2000 Santa Cruz, SC-390124), VCP (1:20000, K331).

**15. Lentivirus generation**

Lentiviral vectors were generated by transfecting HEK293T cells with lentivirus packaging plasmids (pMDLg/pRRE, VsVG and pRSV-REV) with the desired vectors as previously described (Pang et al., 2011) using lipofectamine 3000. The following plasmids were used: pMDLg/pRRE (Addgene 12251), pRSV-Rev (Addgene #12253), pCMV-VSV-G (Addgene #8454), FUW-M2rtTA (Addgene #20342), FUW-TetO-Ngn2-P2A-puromycin (Addgene #52047), FUW-TetO-Ascl1-T2A-puromycin (Addgene #97329), FUW-TetO-Dlx2-IRES-hygromycin (Addgene #97330), TdTomato (Addgene #197033). Lentiviral particles were collected in mTESR+ media and stored at -80°C until further use.

**16. Neuron differentiation and coculture**

hiPSCs were dissociated with Accutase (innovative cell technologies AT-104), cells were counted and $2\times10^5$ cells were plated per well in 6-well plates coated with Matrigel (Cornig 354234) in mTESR+ (Stem cell technology 100-0275) with CETP cocktail (Chroman 1 50nM MedChemExpress HY-15392, Emricasan 5mM Selleck chem S7775, Polyamine supplement 1× SigmaAldrich P8483, trans-ISRIB 700nM R&D systems 5284). A mixture of virus was added to the cell media before plating: i) Ngn2 + rtTA was added for excitatory neuron differentiation (Zhang et al., 2013), ii) Ascl1 + Dlx2 + rtTA was added for inhibitory neuron differentiation (Yang et al., 2017). Excitatory neurons were also transduced with a lentivirus with a plasmid expressing TdTomato on day 4. to distinguish them from inhibitory neurons. On day 1, the media was changed to Neurobasal (Gibco 21103-049) supplemented with B27 (Gibco 17504044) and glutaMAX (Gibco 35050061), doxycycline (2µg/ml, MP biomedicals 198955) was added to the media and kept for 7 days. On day 2 and 3 infected cells were selected with Puromycin (1µg/ml, Sigma-Aldrich P8833) for excitatory neurons, or Puromycin (1µg/ml) and Hygromycin (100µg/ml, Sigma-Aldrich H9773) for inhibitory neurons. On day 4, $8\times10^3$ primary mouse glia were plated into Matrigel coated wells in a 96 well plate. On day 5 induced neurons were dissociated with Accutase and counted, $12\times10^3$ excitatory and $6\times10^3$ inhibitory cells were seeded per well into the coverslips with mouse glia in neurobasal media with 5% FBS (R&D systems S11550) and CEPT cocktail. On day 6 media was changed with neurobasal (with B27 and GlutaMAX) 5% FBS with BDNF (10ng/ml, PeproTech 10781-164), GDNF (10ng/ml, PeproTech 10781-226) and NT3 (10ng/ml, PeproTech 10781-174), Cytosine β-D-arabinofuranoside (AraC 2-4µM Sigma-Aldrich C1768) was added to the media to stop glia proliferation. Half the media was changed every 5 days with neurobasal 5% FBS with BDNF, GDNF and NT3. On day 35 cells were washed 2 times with PBS 1× and fixed with 4% PFA for 30 min. Cells were left in PBS 0.02% sodium azide until staining.

**17. High-content imaging**

For immunofluorescence staining, **h**iPSC-derived neurons were washed twice with 1× PBS and fixed with 4% PFA for 30 min in a 96 well optical bottom plate with a polymer base (Fisher Scientific: 12-566-70) at Rutgers University (New Brunswick, NJ). Fixed neurons were stored at 4ºC in 1× PBS with 0.02% sodium azide and shipped overnight to NorthShore research Institute (Evanston, IL). Neurons were permeabilized in 1× PBS with 0.5% Triton X-100 for 15 minutes at RT without shaking. After blocking with 3% BSA and 0.1% Triton X-100 in 1× PBS for 1hr at RT, the neurons were stained with primary antibodies, mouse anti-Synapsin 1 (1:500), goat anti-tdTomato (1µg/ml), and chicken anti-MAP2 (1:5000), in blocking buffer for 1.5hr at RT. The samples were washed three times in 1× PBS with 0.1% Triton X-100 (0.1% PBST) for 5 min each, and incubated with the secondary antibodies Donkey anti-mouse Alexa 488 (1:1000), donkey anti-goat Alexa 568 (1:1000), and donkey anti-chicken Alexa 647 (1:1000) in blocking buffer for 1hr at RT in the dark. Next, the neurons were washed twice with 0.1% PBST for 5 min, and incubated with DAPI (0.5µg/mL, Fisher Scientific, EN62248) at RT for 10 min. Neurons were washed with 0.05% sodium azide in PBS. The plate was stored at 4℃ and allowed to warm to RT before imaging.

For Image acquisition, the neurons were imaged using Molecular Devices (San Jose, CA) ImageXpress Micro Confocal High-Content Imaging System at both 20× and 40×. The laser wavelengths used were DAPI, FITC, Texas Red, and Cy5. Each well in the 96 well plate was imaged at 8 sites for 40× and 9 sites for 20× with 8-10 *z* stacks at 1µm step size. For the 40× objective the pixel size is 0.3438µm$^2$ with a pinhole of 60µm, 20× objective pixel size is 0.6842 µm$^2$ also with a pinhole of 60µm.

For image analyses, the acquired mages were analyzed as 2D maximum projection. The first two morphometrics, the mean number of neurite branches per cell and the mean length of neurite outgrowth per cell, were analyzed with the built-in Neurite Outgrowth Application Module within the MetaXPress 6 software, version 6.7.2.290. Both mean number of neurite branches per cell and the mean length of neurite outgrowth per cell were calculated based on DAPI stain as nuclear marker and tdTomato stain, which labels excitatory neurons (see generation of neuron culture methods), as neurite and cell body marker. Cell bodies were defined with approximate maximum width of 30µm, a minimum area of 300µm$^2$, and a pixel value of at least 1500 above local background level. Nuclei were identified with approximate minimum width of 8µm, an approximate maximum width of 20µm, and a pixel value of at least 1500 above local background level. Neurite outgrowths were determined with a maximum width of 2µm, minimum projection length of 15µm from the cell body, and a pixel value of at least 500 above local background level. The 20× objective images were used for neurite outgrowth analysis. For assaying the third morphometrics, excitatory synapse density, we used an in-house generated custom synaptic assay module with MetaXPress 6 software. In specific, puncta were identified through Synapsin1 staining with an approximate minimum width of 0.5µm, an approximate maximum width of 2µm, and a minimum pixel value of 2500 above local background level. The number and area of Synapsin1 positive puncta within the colocalized MAP2 and tdTomato signals were used for analysis. The puncta density was generated by the number or total area of puncta within the colocalized MAP2 and tdTomato staining divided by the area of MAP2+& tdTomato+ signal within the neurites. The 40× objective images were used for synaptic puncta density analysis.

##### 18. Statistical analyses

Student's *t*-test was used for comparing statistical differences between two groups, with at least 3 biological replicates. Nominal *P* values were presented. Pearson's correlation was used to evaluate the correlations between two groups. For high content imaging, the cellular phenotypic measurements were from 8 replicates (wells with different cell cultures) and the data of each well were derived from 9 images. For determining chromosomal abnormalities in SNP e-Karyotyping, we use the statistical cut-off as described in the method (Weissbein *et al.*, 2016).
